# CrRLK1L receptor-like kinases HERCULES RECEPTOR KINASE 1 and ANJEA are female determinants of pollen tube reception

**DOI:** 10.1101/428854

**Authors:** Sergio Galindo-Trigo, Noel Blanco-Touriñán, Thomas A. DeFalco, Eloise S. Wells, Julie E Gray, Cyril Zipfel, Lisa M Smith

## Abstract

Communication between the gametophytes is vital for angiosperm fertilisation. Multiple CrRLK1L-type receptor kinases prevent premature pollen tube burst, while another CrRLK1L protein, FERONIA (FER), is required for pollen tube burst in the female gametophyte. We report here the identification of two additional CrRLK1L homologues, HERCULES RECEPTOR KINASE 1 (HERK1) and ANJEA (ANJ), which act redundantly to promote pollen tube burst at the synergid cells. HERK1 and ANJ localise to the filiform apparatus of the synergid cells in unfertilised ovules, and in *herk1 anj* mutants a majority of ovules remain unfertilised due to pollen tube overgrowth, together indicating that HERK1 and ANJ act as female determinants for fertilisation. As in *fer* mutants, the synergid cell-specific, endomembrane protein NORTIA (NTA) is not relocalised after pollen tube reception; however, unlike *fer* mutants, reactive oxygen species levels are unaffected in *herk1 anj* double mutants. Both ANJ and HERK1 associate with FER and its proposed co-receptor LORELEI (LRE) *in planta*. Together, our data indicate that HERK1 and ANJ act with FER to mediate female-male gametophyte interactions during plant fertilisation.

## Introduction

Fertilisation is a critical point in the life cycle of any sexually reproducing organism. In flowering plants, gametes are enclosed in gametophytes, multicellular structures that develop in the reproductive organs of the flower. The pollen grain constitutes the male gametophyte, with each grain generating a pollen tube in the form of a rapidly growing cellular protrusion that delivers the male gametes, or sperm cells, through the style tissues into the ovule. Female gametophytes develop inside the ovule and contain the female gametes within an embryo sac; the egg cell and central cell. The process of double fertilisation in angiosperms consists of the fusion of a sperm cell with each of the female gametes. If fertilisation is successful, the embryo and endosperm develop from the egg cell and central cell fertilisations, respectively. For double fertilisation to occur, the male and female gametophytes must engage in a molecular dialog that controls pollen tube attraction towards the ovule entrance, or micropyle, the arrest of pollen tube growth and the release of the sperm cells in the correct location within the ovule (see (Dresselhaus et al, 2016) for a detailed review).

The synergid cells occupy the micropylar portion of the female gametophyte, and aid communication between the gametophytes. As such, their cytoplasm is densely occupied by endomembrane compartments, reflective of a highly active secretion system generating messenger molecules (Higashiyama, 2002). The filiform apparatus appears at the outermost pole, a thickened and intricate cell wall structure that represents the first contact point between female and male gametophytes prior to fertilisation (Huang & Russell, 1992). Synergid cells secrete small cysteine-rich LURE peptides to guide pollen tubes towards the embryo sac (Okuda et al, 2009). LURE peptides are sensed by two pairs of pollen-specific receptor-like kinases (RLKs), MALE DISCOVERER 1 (MDIS1) and MDIS1-INTERACTING RLK 1 (MIK1), and POLLEN-SPECIFIC RECEPTOR KINASE 6 (PRK6) and PRK3 in Arabidopsis (Takeuchi & Higashiyama, 2016; Wang et al, 2016). These RLKs bind LURE peptides through their extracellular domains at the growing tip of the pollen tubes, triggering directional growth towards the synergid cells (Takeuchi & Higashiyama, 2016; Wang et al, 2016; Zhang et al, 2017).

Within the expanded family of RLKs in Arabidopsis, the *Catharanthus roseus* RLK1-like (CrRLK1L) subfamily has been demonstrated to play several roles during fertilisation. Two pairs of functionally redundant CrRLK1Ls are integral in controlling pollen tip growth, ANXUR1 and 2 (ANX1/2), and BUDDHA’S PAPER SEAL 1 and 2 (BUPS1/2), heterodimerise and ensure pollen tube growth by sensing of two autocrine secreted peptides belonging to the RAPID ALKALINIZATION FACTOR (RALF) family, RALF4 and RALF19 (Boisson-Dernier et al, 2009; Ge et al, 2017; Miyazaki et al, 2009) (Ge et al, 2017; Mecchia et al, 2017). A fifth CrRLK1L protein, ERULUS (ERU), has also been implicated in male-determined pollen tube growth via regulation of Ca^2+^ oscillations (Schoenaers et al, 2017). The CrRLK1L protein FERONIA (FER) accumulates in the filiform apparatus of the synergids where it functions as a female determinant of pollen tube burst and subsequent sperm cell release (Escobar-Restrepo et al, 2007; Huck et al, 2003). Although no extracellular ligand has been identified for FER in a reproductive context, there is evidence for FER activation of a synergid-specific signalling cascade upon pollen tube arrival. This signalling pathway involves the glycosyl-phosphatidylinositol (GPI)-anchored protein LORELEI (LRE) (Li et al, 2015), activation of NADPH oxidases to generate reactive oxygen species (ROS) in the micropyle (Duan et al, 2014), generation of specific Ca^2+^ signatures in the synergid cytoplasm (Ngo et al, 2014), and relocalisation of the Mildew resistance locus O (MLO)-like NORTIA (NTA), an endomembrane compartment protein that affects pollen tube-induced Ca^2+^ signatures in the synergids (Jones & Kessler, 2017; Kessler et al, 2010; Ngo et al, 2014).

Many questions remain about the nature of the communication between gametophytes that controls sperm cell release, and CrRLK1Ls FER, ANX1/2 and BUPS1/2 are potential receptor candidates to mediate this dialog. Here we report the characterisation of CrRLK1Ls HERCULES RECEPTOR KINASE 1 (HERK1) and ANJEA (AT5G59700; ANJ) as female determinants of pollen tube reception in Arabidopsis. We show that HERK1 and ANJ act redundantly at the filiform apparatus of the synergids to control pollen tube growth arrest and burst, representing two new mediators of gametophytic communication and therefore expanding the female-specific toolbox required for fertilisation.

## Results

### HERK1 and ANJEA function redundantly in seed set

To test whether additional Arabidopsis CrRLK1L proteins are involved in reproduction, we obtained T-DNA insertion lines for all seventeen family members. Presence of a homozygous insertion was verified for ten CrRLK1L genes. These verified lines were crossed and double homozygous plants selected in the F2 generation by PCR genotyping (Figure S1A-B for T-DNA lines used further in this study). Stable double homozygous lines were examined for fertility. Through this screen, we identified that double mutants in *HERCULES RECEPTOR KINASE 1 (HERK1)* and *AT5G59700* (hereafter referred to as *ANJEA/ANJ*) have high rates of unfertilised ovules or seeds that abort very early in development, and shorter siliques (Figure 1A). HERK1 and ANJEA are close homologues within the CrRLK1L family (Galindo-Trigo et al, 2016), with 75% identity and 86% similarity at the amino acid level. Loss of *ANJ* gene expression in the double homozygous *herk1-1 anj-1* T-DNA line (hereafter referred to as *herk1 anj*) was confirmed by RT-PCR (Figure S1C), with the *herk1-1* T-DNA insertion previously established to knockout gene expression (Guo et al, 2009a).

**Figure 1.**
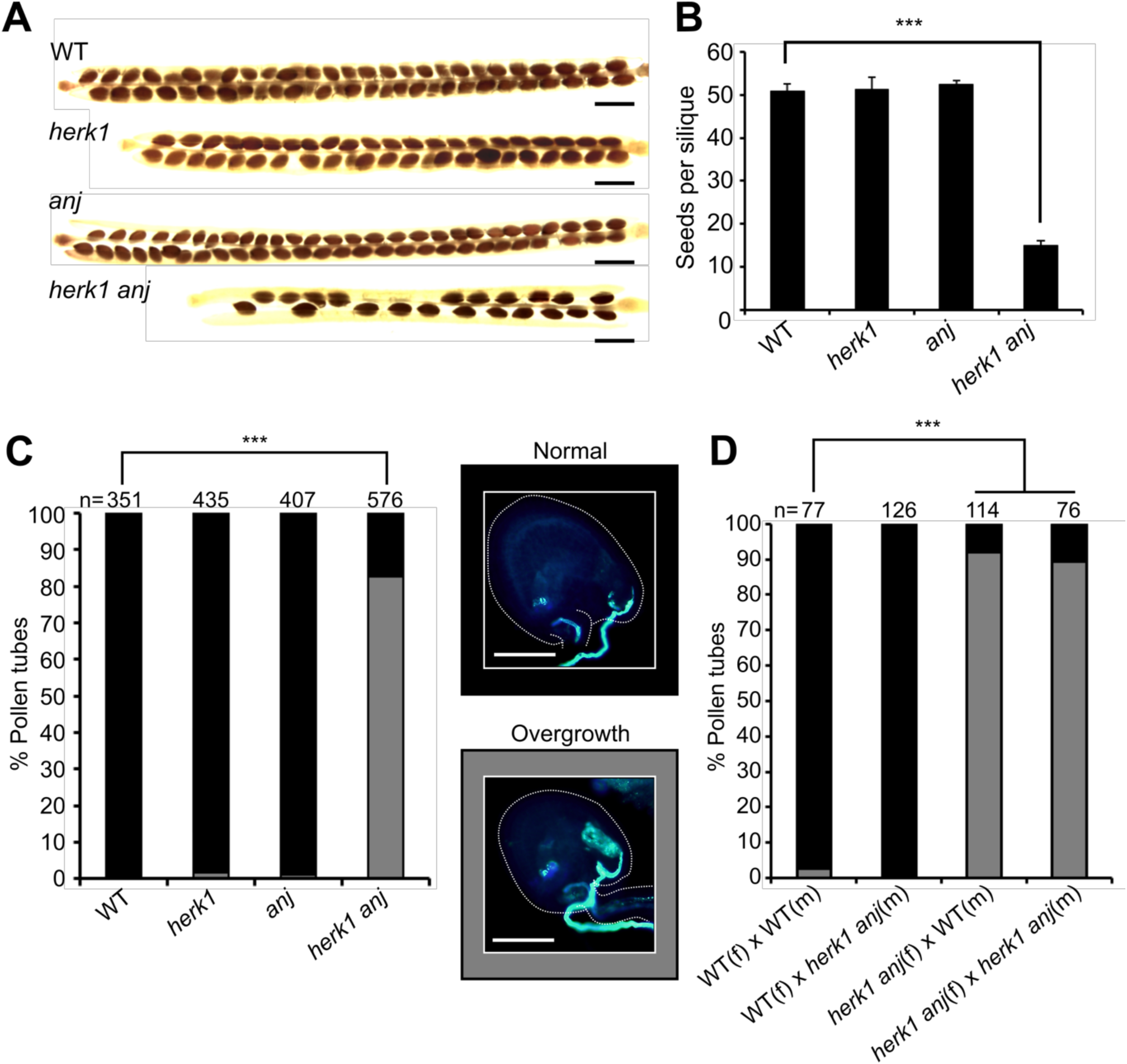
The *herk1 anj* fertility defect is caused by maternally-mediated pollen tube overgrowth. (A) Representative mature siliques from wild-type (WT; Col-0), *herk1*, *anj* and *herk1 anj* plants. Siliques were collected prior to dehiscence and cleared in 0.4M NaOH, 1% Triton X-100. Scale bar = 1 mm. (B) Developing seeds per silique in wild-type, *herk1*, *anj* and *herk1 anj* plants. Fully expanded siliques were dissected and photographed under a stereomicroscope. *n* = 15 (four independent experiments with at least three plants per line and five siliques per plant). Data presented are means ± SEM. *** p<0.001 (Student’s *t*-test). (C) Percentage of pollen tubes with normal reception at the female gametophyte (black bars) and with overgrowth (grey bars) as assessed by aniline blue staining. 15 self-pollinated stage 16 flowers from wild-type, *herk1*, *anj* and *herk1 anj* were analysed. Legend scale bars = 50 µm. *** p<0.001 (χ-square tests). (D) Aniline blue staining of pollen tube reception in reciprocal crosses between wild-type and *herk1 anj* plants with at least two siliques per cross. Legend as per (C). *** p<0.001 (χ-square tests).

To verify that the low rate of seed set results from functional redundancy between *HERK1* and *ANJ*, we examined seed development in dissected siliques of wild-type, *herk1*, *anj* and *herk1 anj* plants grown in parallel. While single mutants *herk1* and *anj* did not have elevated numbers of unfertilised/aborted seeds compared to wild-type, a high proportion of ovules in *herk1 anj* siliques had not developed into mature seeds, leading to a reduced number of seeds per silique (Figure 1B). We therefore concluded that there is functional redundancy between the HERK1 and ANJ proteins during fertilisation or early seed development.

HERK1 has previously been described to influence cell elongation in vegetative tissues with THESEUS1 and HERK2, with the *herk1 the1-4* and *herk1 herk2 the1-4* mutants displaying a short petiole phenotype, similarly to *fer* mutants (Guo et al, 2009a; Guo et al, 2009b). We further examined the *herk1 anj* mutants for developmental defects in vegetative and reproductive growth, finding no further developmental aberrations (Figure S2A-G). Thus, HERK1 and ANJ do not act redundantly during vegetative growth.

### HERK1 and ANJEA are female determinants of pollen tube burst

Previous studies of CrRLK1L proteins where mutation results in low or absent seed set have identified functions in pollen tube growth (ANX1, ANX2, BUPS1, BUPS2 and ERU; (Boisson-Dernier et al, 2009; Ge et al, 2017; Mecchia et al, 2017; Miyazaki et al, 2009; Schoenaers et al, 2017)) and female-mediated pollen tube burst at the synergids (FER (Escobar-Restrepo et al, 2007)). To test which step in fertilisation is impaired in the *herk1 anj* mutant, we tracked pollen tube growth through the style in single and double mutants. In all plant lines, aniline blue staining revealed that the pollen tubes targeted the female gametophytes correctly (Figure S3). However, closer examination of the ovules revealed pollen tube overgrowth at high frequency in *herk1 anj* mutants. While pollen tube overgrowth is rare in wild-type and single mutants, 83% of pollen tubes failed to burst upon entering ovules in the double mutant (Figure 1C). The 83% of ovules exhibiting pollen tube overgrowth is notably higher than the 71% of ovules that fail to develop into seeds (Figure 1B,C), indicating that in some cases fertilisation occurs in the presence of pollen tube overgrowth.

In *fer* mutants, pollen tube overgrowth occurs due to maternal defects in male-female gametophyte communications (Duan et al, 2014; Escobar-Restrepo et al, 2007; Huck et al, 2003). To confirm that HERK1 and ANJ are female determinants of pollen tube burst, we performed reciprocal crosses between the *herk1 anj* mutant and wild-type plants, as well as control crosses within each plant line. While wild-type Col-0 (female; f) x *herk1 anj* (male; m) crosses resulted in 1% of ovules with pollen tube overgrowth, over 90% of pollen tubes exhibited overgrowth in *herk1 anj* (f) x wild-type (m) crosses, indicating that pollen tube overgrowth is a maternally-derived phenotype in *herk1 anj* mutants (Figure 1D). As expected, pollen tube overgrowth was observed in only 3% of the ovules in the control wild-type (f) x wild-type (m) crosses, while 89% of ovules had overgrowth of the pollen tube in *herk1 anj* (f) x *herk1 anj* (m) crosses.

To verify that the reproductive defect is due to the disruption of the *HERK1* and *ANJ* genes and does not arise from additional T-DNA insertions, we reintroduced the *HERK1* and *ANJ* genes into the *herk1 anj* background to test for complementation of the pollen tube overgrowth phenotype. We generated *pHERK1::HERK1* and *pANJ::ANJ-GFP* constructs and obtained *pFER::HERK1-GFP* (Kessler et al, 2015). A *pBRI1::HERK1-GFP* construct has previously been used to complement the *herk1* mutant (Guo et al, 2009a), and we found that while *pHERK1::HERK1* could be generated, *pHERK1::HERK1-GFP* could not be cloned due to toxicity in several bacterial strains. In the developing ovules of five independent T1 plants where a hemizygous insertion would segregate 50:50, expression of *pFER::HERK1-GFP* or *pANJ::ANJ-GFP* constructs in the *herk1 anj* background reduced pollen tube overgrowth by ~50%, as did a *pHERK1::HERK1* construct (Figure S4A). Complementation indicates that these reporter constructs produce functional proteins and confirms that the T-DNA insertions in the *HERK1* and *ANJ* genes are responsible for pollen tube overgrowth. We conclude that HERK1 and ANJ are female determinants of pollen tube burst and therefore named *AT5G59700* after a fertility goddess in Australian aboriginal mythology, Anjea.

The kinase activity of FER is not required for its control of pollen tube reception in ovules (Kessler et al, 2015). We therefore tested for complementation of the *herk1 anj* reproductive defect with kinase-dead (KD) versions of HERK1 and ANJ generated by targeted mutagenesis of key residues within the kinase activation loop (D609N/K611R for HERK1 and D606N/K608R for ANJ; (Knighton et al, 1991)). *pHERK1::HERK1-KD* and *pANJ::ANJ-KD-GFP* were also able to complement the pollen overgrowth phenotype, indicating that the kinase activity of these RLKs is not required for their function in fertilisation (Figure S4B). The similarity in the mutant phenotypes, cellular localisation and the dispensable kinase activity in HERK1/ANJ and FER suggests they may act in the same signalling pathway as co-receptors or as parallel receptor systems.

### HERK1 and ANJEA are localised to the filiform apparatus

To explore the localisation of HERK1 and ANJ in the female gametophyte and hence gain insight into the possible function of HERK1/ANJ in fertilisation, we made *promoter::H2B-TdTomato* transcriptional fusions where expression of either the *HERK1* or *ANJ* promoter should direct nuclear localisation of an RFP signal. Both *HERK1* and *ANJ* were strongly expressed in unfertilized embryo sacs, with expression of *HERK1* in the two synergid cells, egg cell and central cell of 4-cell stage female gametophytes and *ANJ* expression restricted to the two synergid cells (Figure 2A-D). As HERK1 and ANJ must be expressed in the same cells for a genetic interaction to occur, this restricts their potential function in the female gametophyte during fertilisation to the synergid cells.

**Figure 2.**
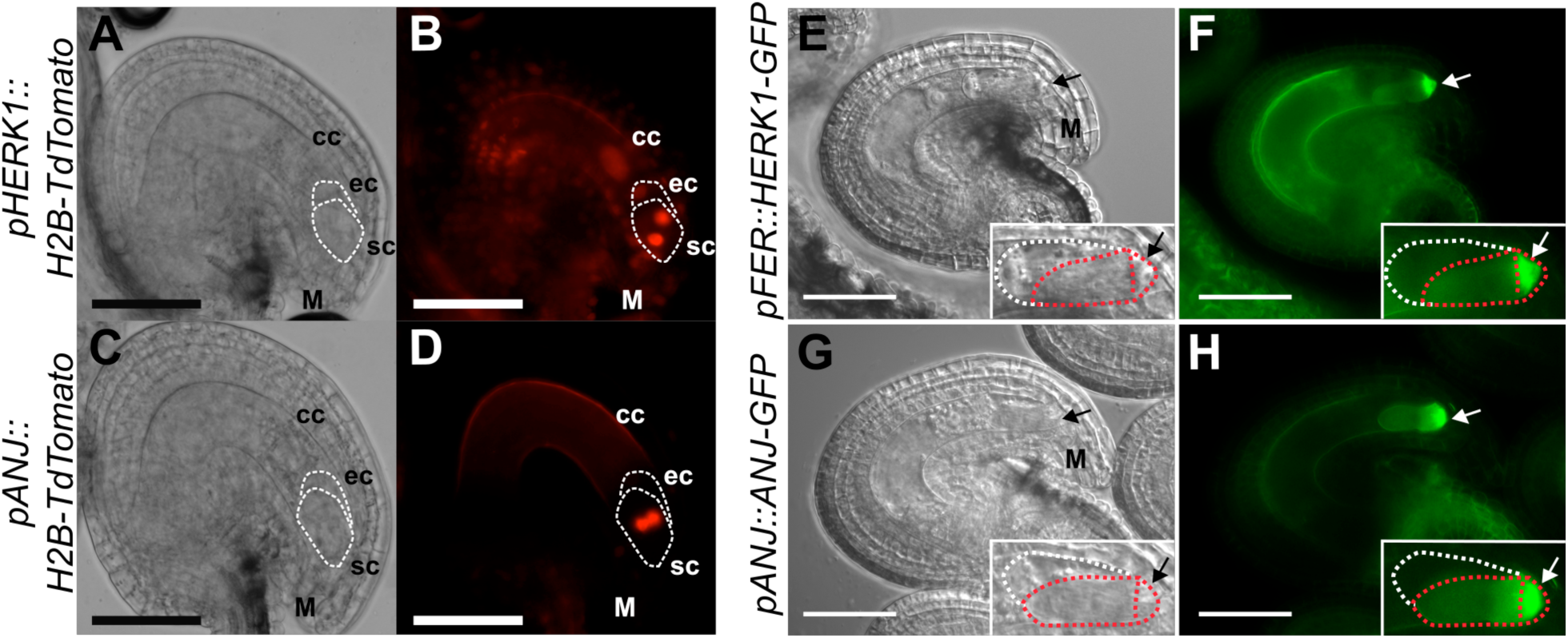
HERK1 and ANJ are expressed in the female gametophyte and localise to the filiform apparatus of the synergid cells. (A-B) Expression of *pHERK1::H2B-TdTomato* in mature ovules. (C-D) Expression of *pANJ::H2B-TdTomato* in mature ovules. Scale bars = 50 µm. White dotted lines delineate the egg cell and synergid cells. (E,F) Localisation of HERK1-GFP in the synergid cell from the *pFER::HERK1-GFP* construct in (F) and corresponding differential interference contrast (DIC) image in (E). (G,H) Localisation of ANJ-GFP in the synergid cell from the *pANJ::ANJ-GFP* construct in (H) and corresponding DIC image in (G). White and red dotted lines delineate the egg cell and synergid cells, respectively. Scale bars = 50 µm. M, micropyle. Arrows, filiform apparatus.

We next generated *promoter::GUS* (β-glucuronidase) transcriptional fusions to gain insight into the expression of these genes at a tissue level. *pHERK1::GUS* is also expressed in the style, ovary walls and stamens (Figure S5A-D), whereas *pANJ::GUS* expression is detected in stigmas and stamens (Figure S5E-H). No expression was detected in pollen grains within mature anthers, although HERK1 was expressed in some developing pollen grains (Figure S5B,D,H). Thus *HERK1* and *ANJ* are expressed in multiple reproductive tissues, with the pattern of expression suggesting the fertilisation defect may arise through a biological function in the junction of the stigma and style, or in the female gametophyte where *HERK1* and *ANJ* gene expression overlaps in the synergid cells.

To further examine HERK1 and ANJ expression and subcellular localisation in ovules, we used the *pANJ::ANJ-GFP* and *pFER::HERK1-GFP* constructs that complement the fertilisation phenotype. Examination of fluorescent signals from HERK1-GFP and ANJ-GFP fusion proteins in the female gametophyte showed that they were strongly localised to the filiform apparatus of the synergid cells (Figure 2E-H). The filiform apparatus is a structure formed by dense folds in the plasma membrane and cell wall where the regulators of fertilisation FER and LRE also localise (Capron et al, 2008; Escobar-Restrepo et al, 2007; Tsukamoto et al, 2010b). This specific cellular localisation supports the hypothesis that HERK1 and ANJ could function in the same pathway as FER and LRE. While loss of FER or LRE alone leads to a reproductive defect caused by pollen tube overgrowth in the ovule (Capron et al, 2008; Escobar-Restrepo et al, 2007), HERK1 and ANJ are functionally redundant, such that HERK1 and ANJ could act as alternative co-receptors for FER and/or LRE during male-female interactions.

### NORTIA relocalisation after fertilisation is impaired in *herk1 anj* mutants

Previous reports point to an interdependence between FER, LRE and NTA in their respective cellular localisations (Kessler et al, 2010; Li et al, 2015). FER only accumulates in the filiform apparatus if functional LRE is present, and NTA relocalisation towards the filiform apparatus upon pollen tube arrival is dependent on FER (Kessler et al, 2010; Li et al, 2015). As HERK1 and ANJ may act in the same signalling pathway as FER, we tested the localisation of fluorescence-tagged HERK1, ANJ, FER, LRE and NTA in the *herk1 anj* and *lre-5* backgrounds (Figure 3A). Localisation within the synergids of FER-GFP, LRE-Citrine and NTA-GFP was not affected by *herk1 anj* mutations. Similarly, HERK1-GFP and ANJ-GFP localised to the filiform apparatus in the *lre-5* background. Contrary to previous findings (Li et al, 2015), under our conditions FER-GFP accumulation in the filiform apparatus was not impaired in *lre-5* plants (n>25; FER-GFP was found at the filiform apparatus in all ovules checked). To verify that FER subcellular localisation was not affected in *lre-5* under our growth conditions, we quantified the mean fluorescence intensity across the filiform apparatus (FA) and synergid cytoplasm (SC) to calculate the ratio of FA:SC fluorescence intensity (Figure S6A). When compared across the wild-type, *herk1 anj* and *lre-5* genotypes, the mean FA:SC fluorescence intensity ratios were not significantly different, indicating no effect on FER-GFP localisation to the FA in plants lacking LRE or HERK1/ANJ. Furthermore, we found no differences in the percentage of ovules presenting moderate or severe mislocalisation of FER-GFP in the synergid cells in wild-type, *herk1 anj* or *lre-5* plants (Student’s *t* tests, p>0.05; Figure S6B). Therefore, we found no dependency on HERK1/ANJ or LRE for localisation of FER, LRE, HERK1, ANJ or NTA within the synergids of unfertilised ovules.

**Figure 3.**
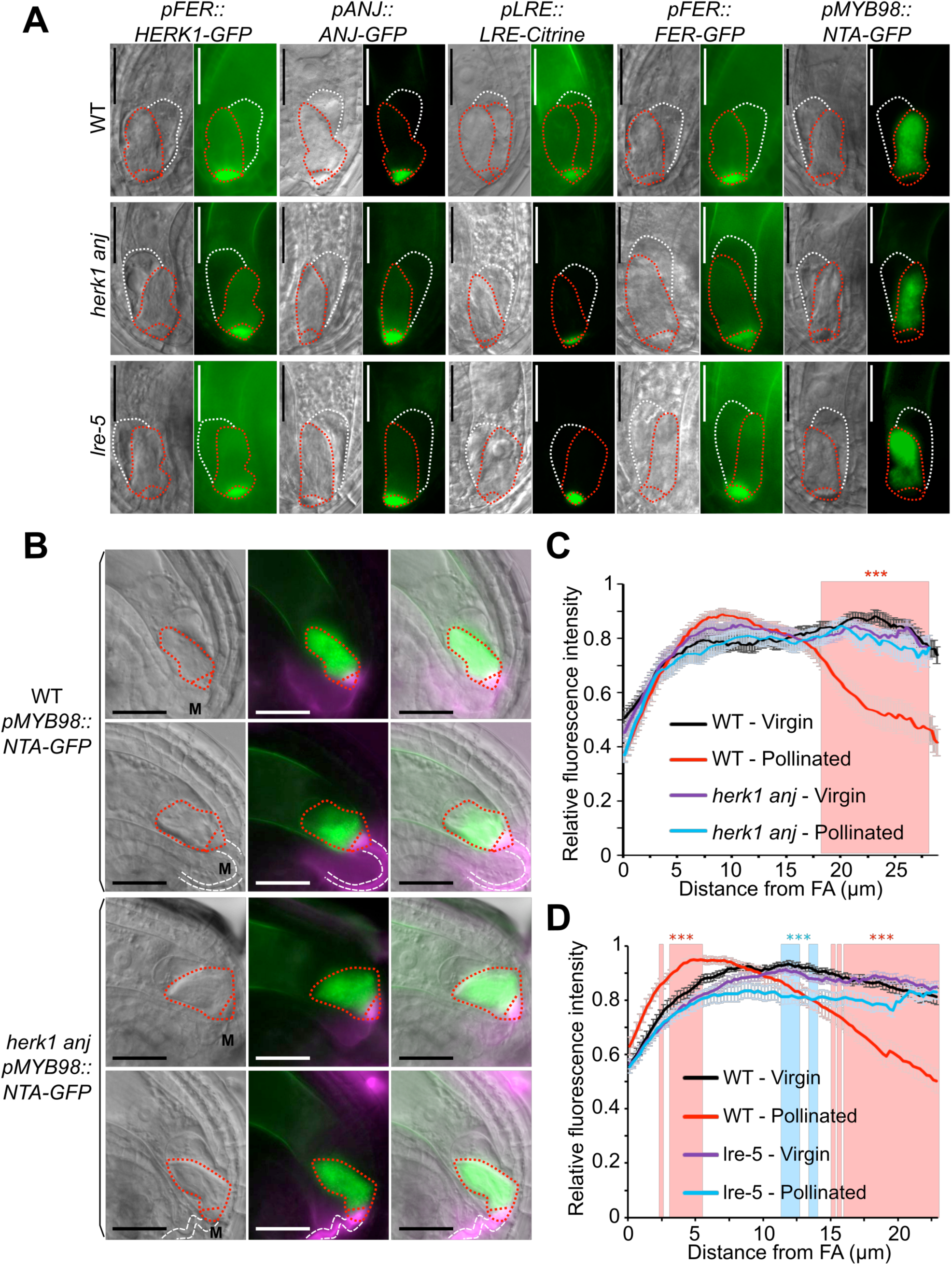
Normal synergid localisation of HERK1, ANJ, LRE, FER and NTA pre-fertilisation and impaired relocalisation of NTA after pollen tube reception in *herk1 anj* and *lre-5*. (A) Localisation of HERK1, ANJ, LRE, FER and NTA in the synergid cell of wild-type (Col-0; WT), *herk1 anj* and *lre-5* in unfertilised ovules, as shown by *pFER::HERK1-GFP*, *pANJ::ANJ-GFP*, *pLRE::LRE-Citrine*, *pFER::FER-GFP* and *pMYB98::NTA-GFP*. DIC and fluorescence images are shown, left to right, respectively. White and red dotted lines delineate the egg cell and synergid cells, respectively. Scale bars = 25 µm. (B) Localisation of NTA in the synergid cell of wild-type and *herk1 anj* plants before (upper panels) and after (lower panels) pollen tube arrival. In green, NTA localisation as shown by *pMYB98::NTA-GFP* fluorescence. In magenta, callose of the filiform apparatus and pollen tube stained with SR2200. From left to right, images shown are DIC, merged fluorescence images, and merged images of DIC and fluorescence. White and red dotted lines delineate the pollen tube and synergid cells, respectively. Scale bars = 25 µm. M, micropyle. (C-D) Profile of relative fluorescence intensity of NTA-GFP along the synergid cells of wild-type and *herk1 anj* ovules (C); and wild-type and *lre-5* ovules (D) before (virgin) and after (pollinated) pollen arrival. Data shown are means ± SEM, n = 25. *** p<0.001 (Student’s *t*-test). FA, filiform apparatus.

To determine whether NTA relocalisation in synergid cells upon pollen tube arrival depends on functional HERK1 and ANJ, we transformed *pMYB98::NTA-GFP* into the *herk1 anj* background. Using SR2200-based callose staining to visualise the filiform apparatus and pollen tube, we observed NTA-GFP fluorescence intensity across the length of the synergid cell. In unfertilised ovules, NTA-GFP fluorescence is evenly distributed across the length of the synergid cell in wild-type and *herk1 anj* plants (Figure 3B). Wild-type fertilised ovules have a shift in the fluorescence intensity pattern, with NTA accumulation towards the micropylar end of the synergid cytoplasm and a decrease in relative fluorescence intensity towards the chalazal end (Figure 3B-C). This response is absent in *herk1 anj* fertilised ovules in which the relative fluorescence intensity pattern is indistinguishable from that of unfertilised ovules, indicating a requirement for HERK1/ANJ in NTA relocalisation upon pollen tube perception.

Whether LRE is dispensable for NTA relocalisation upon pollen tube arrival has not previously been tested. We therefore transformed the *pMYB98::NTA-GFP* construct into the *lre-5* genetic background and repeated the assay above to examine whether LRE is required for NTA relocalisation as for HERK1, ANJ and FER (Kessler et al, 2010). While a region of statistically lower signal intensity was present around the middle of the synergids in pollinated *lre-5* ovules compared to wild-type virgin ovules (Figure 3D), there was no significant shift in signal toward the filiform apparatus upon fertilisation as observed for wild-type pollinated ovules. Therefore, under our growth conditions, NTA relocalisation at pollen tube arrival is also affected by a loss of LRE.

As reported by Ngo and colleagues (2014), the journey of the pollen tube does not conclude upon contact with the filiform apparatus of the synergid cells (Ngo et al, 2014). Pollen tubes transiently arrest growth upon contact with the synergid; they then grow rapidly along the receptive synergid and towards the chalazal end, before burst and release of the sperm cells (Ngo et al, 2014). To observe this process in detail, we used TdTomato-tagged pollen and monitored NTA-GFP localisation at different stages of pollen tube growth within the ovule. The shift in NTA-GFP localisation was noted in ovules in which the pollen tube had grown past the filiform apparatus and ruptured, rather than upon pollen tube arrival at the filiform apparatus (Figure S7A). Interestingly, in rare cases when pollen tube burst occurred normally in the *herk1 anj* background, the fluorescence shift towards the micropyle had also taken place (Figure S7A). In both cases, NTA-GFP did not appear to accumulate in the filiform apparatus (Figure S7B). Our results differ from the interpretation of previous reports that NTA is polarly relocalised from endomembrane compartments to the plasma membrane in the filiform apparatus, instead supporting a more generalised relocalisation within the synergid cytoplasm towards the micropylar end, at least under our growth conditions. We propose that HERK1, ANJ and LRE, similarly to FER, act upstream of NTA relocalisation in the signalling pathway.

### ROS production is not affected in mature *herk1 anj* ovules

ROS levels in *fer-4* and *lre-5* ovules have been reported to be significantly lower than in wild-type with the implication that, as hydroxyl free radicals can induce pollen tube burst (Duan et al, 2014), reduced ROS levels could be responsible for pollen tube overgrowth. To assess whether HERK1 and ANJ also act upstream of ROS accumulation in the ovules, we used H_2_DCF-DA to measure ROS levels on a categorical scale in *herk1 anj*, *lre-5* and *fer-4* ovules (Figure S8A). To ensure that all ovules were fully developed prior to ROS measurement, we emasculated stage 14 flowers and allowed them to develop for a further 20 hours. At 20 hours after emasculation (HAE), all ovules had reached the mature 7-celled or 4-celled pollen-receptive stages in all backgrounds tested (Figure S10B; (Christensen et al, 1998; Yadegari & Drews, 2004)). Across three independent experiments, we confirmed that ROS levels are significantly lower in *fer-4* ovules compared to wild-type (Figure 4B and S8C), indicating the that ROS assay is functional in our hands and able to distinguish changes in ROS levels. However, we found that ROS levels are consistently comparable to wild-type in mature ovules of *herk1 anj* and *lre-5* (Figure 4B and S8C). To verify that the fertilisation defect is not rescued in the *herk1 anj* and *lre-5* genotypes at 20 HAE, we confirmed that pollen tube overgrowth still occurs when ovules are fertilised at this stage (Figure 4C). Taken together, these results suggest that FER acts upstream of ROS accumulation in ovules prior to pollen tube arrival while, under our experimental conditions, HERK1, ANJ and LRE are not required for this process. As these results conflict with a previous study showing lower ROS levels in *lre-5* ovules (Duan et al, 2014), this suggests that the function of LRE in ROS production may be environmentally sensitive. Our results do not preclude that pollen tube arrival-induced ROS signalling in the synergid cells is affected in *herk1 anj* and *lre-5*, however differences in transient synergid-specific ROS burst cannot be quantified in our *in vitro* system.

**Figure 4.**
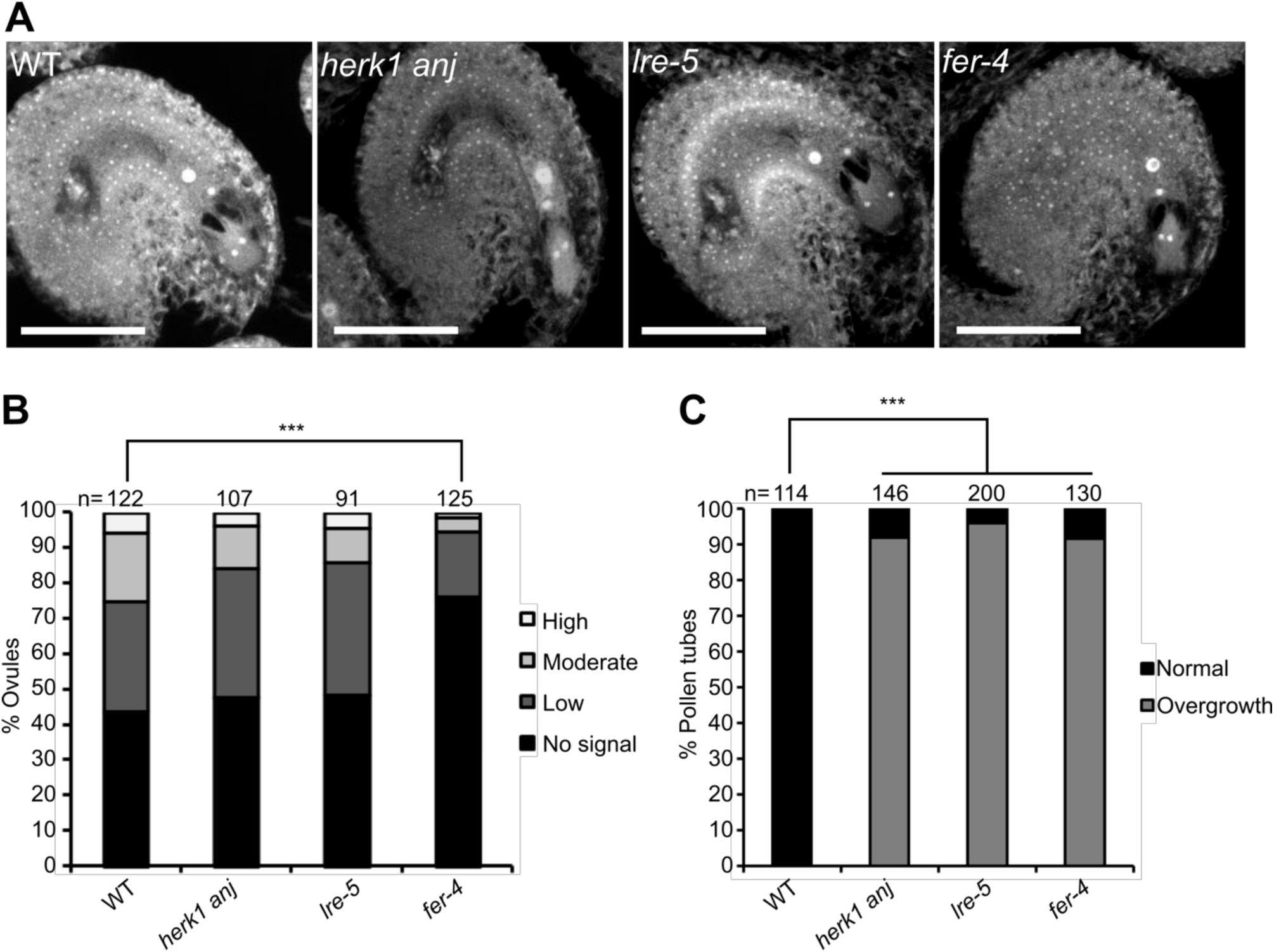
*herk1 anj* mature female gametophytes are morphologically normal and unaffected in ROS production at the micropyle. (A) Representative images of ovules from wild-type (Col-0), *herk1 anj*, *lre-5* and *fer-4* 20 hours after emasculation (HAE) displaying the mature female gametophyte structure. Images presented here are maximum intensity projections from confocal microscopy images across several z-planes of ovules stained as per (Christensen et al, 1997). Scale bars = 50 µm. (B) Quantification of H_2_CDF-DA staining of ROS in ovules from wild-type, *herk1 anj*, *lre-5* and *fer-4* plants at 20 HAE. Categories are listed in the legend (see also Figure S8A). Ovules dissected from at least five siliques per line. *** p<0.001 (χ-square tests). (C) Percentage of pollen tubes with normal reception at the female gametophyte (black bars) and displaying overgrowth (grey bars) in wild-type, *herk1 anj*, *lre-5* and *fer-4* plants, manually selfed at 20 HAE. Fertilisation events counted from at least three siliques per line. *** p<0.001 (Student’s *t*-test).

### HERK1 and ANJEA interact with LORELEI and FERONIA

LRE and its homolog LORELEI-LIKE GPI-ANCHORED PROTEIN 1 (LLG1) physically interact with RLKs FER, FLAGELLIN SENSING 2 (FLS2) and EF-TU RECEPTOR (EFR) (Li et al, 2015; Shen et al, 2017). Mutations in these GPI-anchored proteins and their associated RLKs result in similar phenotypes, with LRE and LLG1 regarded as co-receptors and/or stabilisers of RLK function (Li et al, 2015; Shen et al, 2017). HERK1, ANJ and FER are closely related RLKs and, given the similarities in reproduction defects and sub-cellular localisation in synergid cells (Figure 3A), we hypothesised that HERK1 and ANJ may act in complex with LRE and/or FER at the filiform apparatus. To examine this hypothesis, we used yeast two hybrid assays to test for direct interactions between the extracellular juxtamembrane domains of HERK1, ANJ (HERK1exJM, ANJexJM) and LRE, as well as the complete extracellular domains of HERK1, ANJ and FER (HERK1-ECD, ANJ-ECD and FER-ECD). Interactions between HERK1exJM and ANJexJM with LRE were detected, as were interactions of FER-ECD and HERK1-ECD with FER-ECD, HERK1-ECD and ANJ-ECD, and of ANJ-ECD with FER-ECD and HERK1-ECD, indicative of a possible direct interaction between these four proteins (Figure 5A-B). Interactions were also tested by yeast two hybrid assays between the kinase domains of HERK1, ANJ and FER (HERK1-KIN, ANJ-KIN and FER-KIN) but interaction between these domains was much weaker (Figure S11).

**Figure 5.**
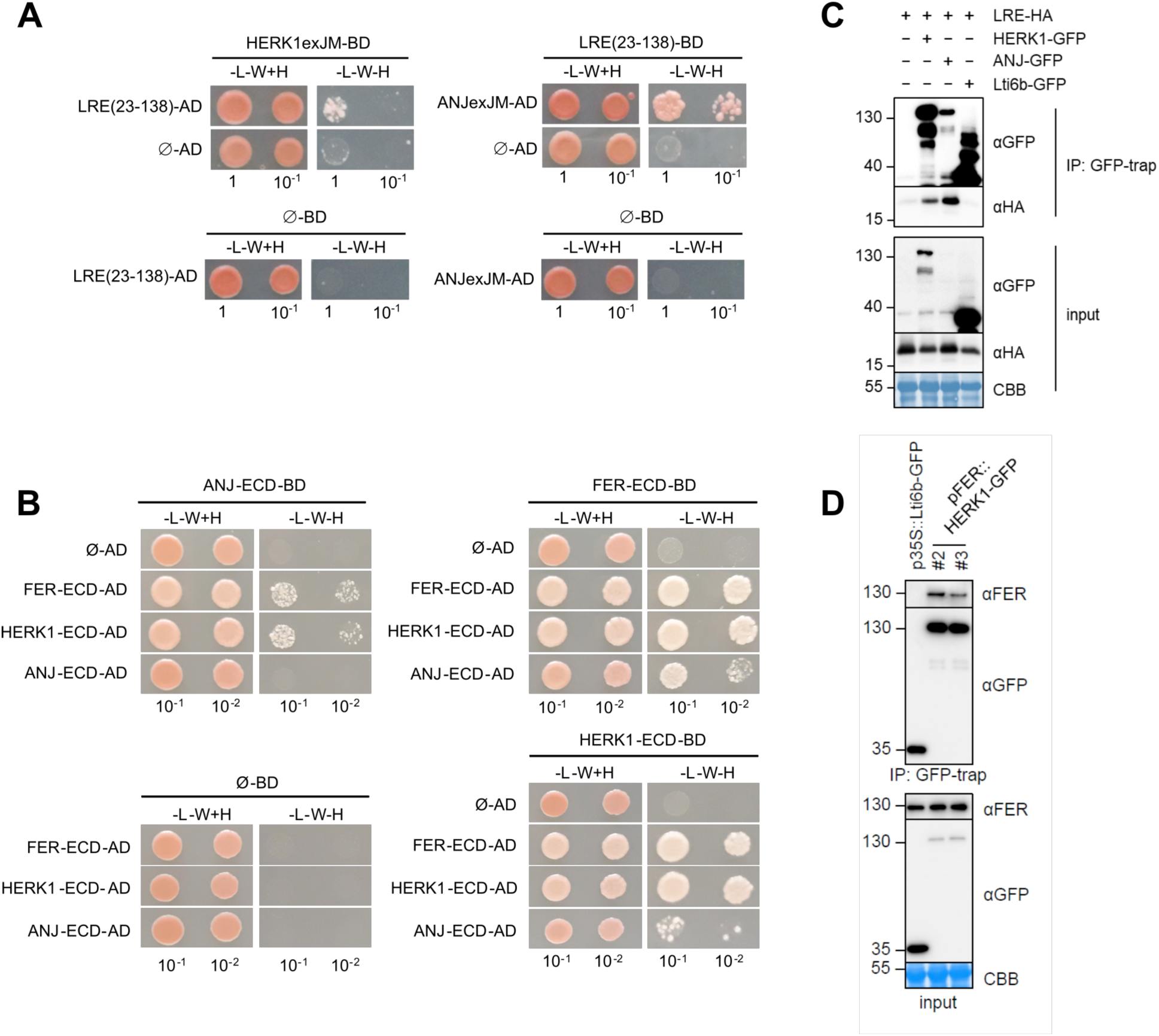
HERK1 and ANJ interact with LRE and FER. (A) Yeast two hybrid (Y2H) assays of the extracellular juxtamembrane domains of HERK1 and ANJ (HERK1exJM and ANJexJM, respectively) with LRE (residues 23-138; signal peptide and C-terminal domains excluded). (B) Y2H assays with the extracellular domains of HERK1, ANJ and FER (HERK1-ECD, ANJ-ECD and FER-ECD, respectively). Ø represents negative controls where no sequence was cloned into the activating domain (AD) or DNA-binding domain (BD) constructs. -L-W-H, growth medium depleted of leucine (-L), tryptophan (-W) and histidine (-H). (C) Co-immunoprecipitation of HA-LRE with HERK1-GFP or ANJ-GFP following 2 days of transient expression in *N. benthamiana* leaves. (D) Co-immunoprecipitation of FER with HERK1-GFP in Arabidopsis seedlings expressing *pFER::HERK1-GFP*. Numbers indicate MW marker sizes in kDa. Assays were performed twice with similar results.

To corroborate interactions of HERK1, ANJ, FER and LRE *in planta*, co-immunoprecipitation assays were performed. In a heterologous system using *Agrobacterium*-mediated transient expression of *pFER::HERK1-GFP, pFER::ANJ-GFP* and *p35S::HA-LRE* in *Nicotiana benthamiana* leaves, HA-LRE co-immunoprecipitated with HERK1-GFP and ANJ-GFP (Figure 5C), confirming that these proteins form complexes *in planta*. Furthermore, *herk1 anj* lines complemented with *pFER::HERK1-GFP* were used to assay the association of HERK1 with endogenous FER using an α-FER antibody (Xiao, Stegmann, Han et al., under revision). FER co-immunoprecipitated with both HERK1-GFP in several independent experiments (Figure 5D), again confirming that these complexes form *in planta*. In an additional genetic approach, we introduced the *lre-5* mutation into the *herk1 anj* background and characterised fertility impairment in triple homozygous *herk1 anj lre-5* plants. No additive effect was observed in the seed set defect in *herk1 anj lre-5* plants compared to *herk1 anj* and *lre-5* mutants (Figure 6A). To test for any additional additive interaction between HERK1, ANJ, FER and LRE at the level of seed set, CRISPR-Cas9 was used with two guide RNAs to generate deletions in *FER* in wild-type, *herk1 anj* and *herk1 anj lre-5* genetic backgrounds. Seed set was analysed in T2 plants grown in parallel with wild-type, *herk1 anj*, *lre-5*, *fer-4* and *herk1 anj lre-5* mutants. No statistically significant difference was found between single, double, triple or quadruple mutants, while all mutants produced significantly fewer seeds than wild-type (Figure S12).

**Figure 6.**
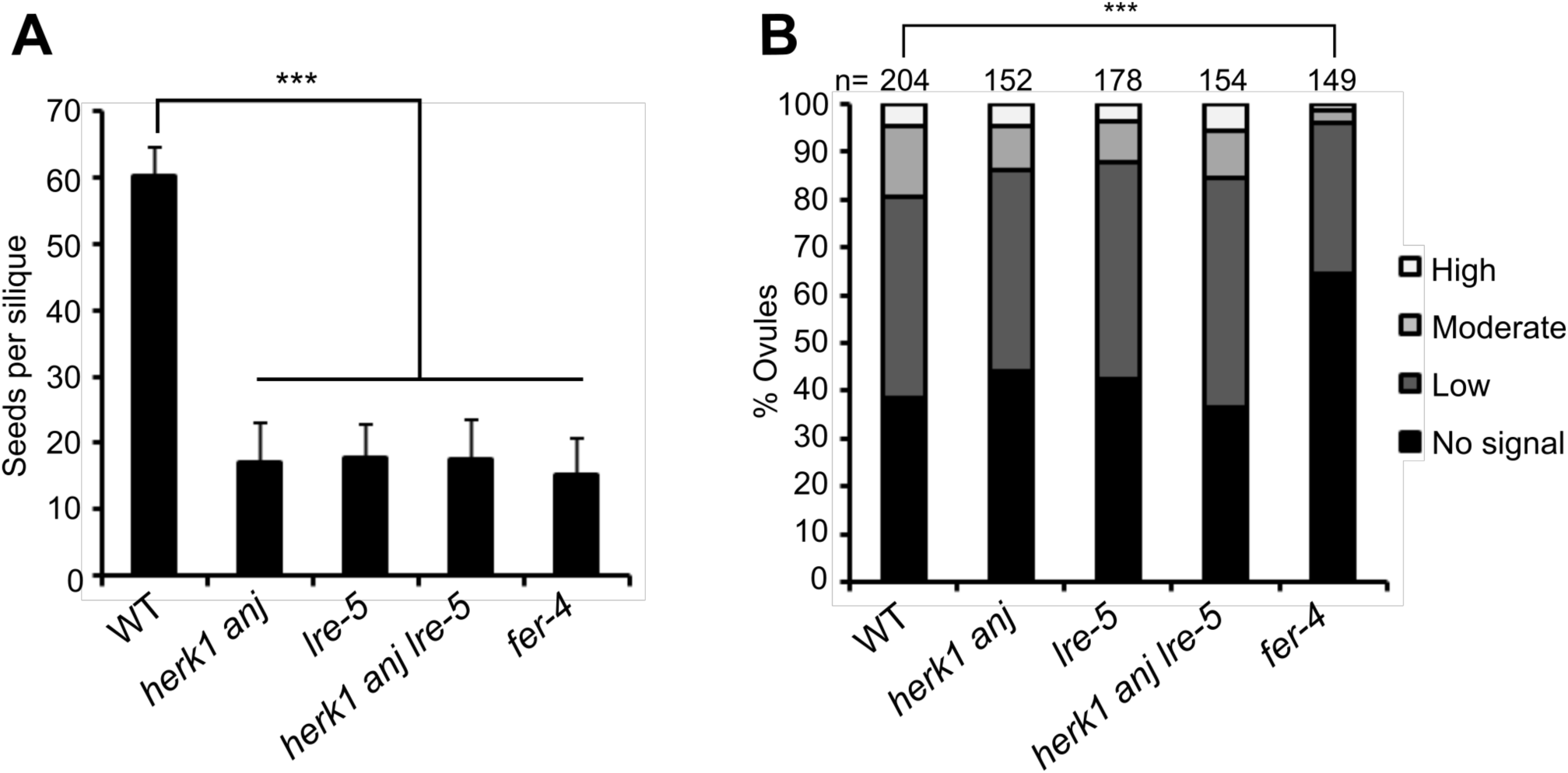
HERK1, ANJ and LRE do not act additively in seed set or ROS production. (A) Quantification of developing seeds per silique in wild-type, *herk1 anj*, *lre-5*, *herk1 anj lre-5* and *fer-4* plants. Fully expanded siliques were dissected and photographed under a stereomicroscope. n = 25. Data presented are means ± SD. *** p<0.001 (Student’s *t*-test). (B) Quantification of the H_2_CDF-DA staining of ROS in ovules from wild-type, *herk1 anj*, *lre-5*, *herk1 anj lre-5* and *fer-4* plants at 20 HAE. Categories are listed in the legend (see also Figure S8A). Ovules dissected from at least five siliques per line. *** p<0.001 (χ-square tests).

It has been reported for several mutations causing pollen tube overgrowth, including *lre* and *fer*, that pollen tube overgrowth is occasionally accompanied by polytubey, where more than one pollen tube enters the ovule (Figure S13A-B; (Capron et al, 2008; Huck et al, 2003)). This is indicative of uninterrupted secretion of attraction signals from the synergid cells, suggesting impaired degeneration of the receptive synergid cell upon pollen tube arrival (Beale et al, 2012; Maruyama et al, 2015). Polytubey has been reported to occur at a rate of ~10% in the progeny of a heterozygous *fer-1* mutant (Huck Dev 2003). To assess whether polytubey occurs in the *herk1 anj* mutant at a similar rate, polytubey was quantified in *herk1 anj* mutants along with *lre-5* and *fer-4* mutants as controls (Figure S13C). Under our growth conditions polytubey was more frequent in *fer-4* mutants (38.6% of fertilized ovules) than previously reported for *fer-1*. Compared to *fer-4, herk1 anj* (24.8% of fertilized ovules) and *lre-5* mutants (27.2% of fertilized ovules) exhibited statistically lower rates of polytubey, whereas *herk1 anj lre-5* mutants presented similar rates to *fer-4* (40.3% of fertilised ovules), indicating that mutations in *HERK1*, *ANJ* and *LRE* may have an additive effect in the attraction of supernumerary pollen tubes.

ROS production in ovules of the triple *herk1 anj lre-5* mutant was measured using H_2_DCF-DA at 20 HAE. In agreement with the seed set phenotype, ROS levels were unaffected in the triple homozygous line (Figure 6B). These results reinforce the hypothesis that HERK1, ANJ and LRE act in the same signalling pathway and, given their cellular localisation and our protein-protein interaction results, we propose that HERK1-LRE-FER and ANJ-LRE-FER form part of a receptor complex in the filiform apparatus of synergid cells which mediates pollen tube reception.

## Discussion

Successful reproduction in angiosperms relies on tightly controlled communication between gametophytes through the exchange of chemical and mechanical cues (Dresselhaus et al, 2016). Here, we describe the role of the RLKs HERK1 and ANJ in early stages of fertilisation in Arabidopsis. HERK1 and ANJ are widely expressed in female reproductive tissues including the synergid cells of ovules, where they are polarly localised to the filiform apparatus. *herk1 anj* plants fail to produce seeds from most ovules due to a maternally-derived pollen tube overgrowth defect. As female gametophytes develop normally in *herk1 anj* mutants, pollen tube overgrowth is likely due to impaired signalling. To clarify the position of HERK1/ANJ in relation to the previously characterised signalling elements of the pollen tube reception pathway, we have shown that NTA relocalisation after pollen tube reception is impaired in *herk1 anj* as described for FER, whereas ROS production at the micropylar entrance of ovules prior to pollen arrival is not affected. Interactions between HERK1/ANJ, FER and LRE lead us to propose receptor complexes containing HERK1-LRE-FER and ANJ-LRE-FER at the filiform apparatus.

Associated with diverse hormonal, developmental and stress responses, FER is regarded as a connective hub of cellular responses through its interactions with multiple partners, including small secreted peptides, cell wall components, other RLKs, GPI-anchored proteins and ROPGEFs (Duan et al, 2010; Feng et al, 2018; Haruta et al, 2014; Hou et al, 2016; Li et al, 2015; Stegmann et al, 2017). As related members of the CrRLK1L family, HERK1 and ANJ have the potential to perform similar roles to FER, and as reported here control pollen tube rupture. Interestingly, control of tip growth in pollen tubes depends on two redundant pairs of CrRLK1Ls; ANX1 and ANX2, and BUPS1 and BUPS2 (Boisson-Dernier et al, 2009; Ge et al, 2017; Mecchia et al, 2017; Miyazaki et al, 2009). ANX1/2 and BUPS1/2 form ANX-BUPS heterodimers to control pollen tube growth by sensing autocrine RALF signals (Ge et al, 2017). In turn, ovular RALF34 efficiently induces pollen tube rupture at the pollen tip, likely through competition with autocrine RALF4/19 (Ge et al, 2017). LEUCINE-RICH REPEAT EXTENSINS (LRXs) constitute an additional layer of regulation during pollen tube growth (Mecchia et al, 2017). LRXs interact physically with RALF4/19 and are thought to facilitate RALF sensing during pollen tube growth (Mecchia et al, 2017; Stegmann & Zipfel, 2017). Here we propose that female control of pollen tube reception is also executed via CrRLK1L heterocomplexes of FER with either HERK1 or ANJ, which could sense pollen tube-derived cues to trigger the female gametophyte to induce pollen tube rupture. Given the multiple CrRLK1L-RALF interactions identified to date (Ge et al, 2017; Gonneau et al, 2018; Haruta et al, 2014; Mecchia et al, 2017), pollen tube-derived RALF signals constitute a potential candidate to induce synergid responses to pollen tube perception. RALF4/19 are continuously secreted at the growing tip of the pollen tube and, while their involvement in pollen growth has been thoroughly studied (Ge et al, 2017; Mecchia et al, 2017), their possible dual role as synergid-signalling activators remains unexplored. Disruption of synergid autocrine RALF signalling upon pollen arrival constitutes another possible model, comparable to that hypothesised for RALF34 and RALF4/19 during pollen growth (Ge et al, 2017). Additionally, LRXs could facilitate RALF perception at the synergid cell to control pollen tube reception.

A second category of putative pollen tube cues involves changes in cell wall properties of the filiform apparatus. As a polarised fast-growing structure, pollen tubes present cell walls that differ from stationary cell types, with particular emphasis on the growing tip where active cell wall remodelling rapidly takes place (Chebli et al, 2012). When the growing tip reaches the filiform apparatus, it temporarily arrests growth, subsequently growing along the receptive synergid cell prior to rupture (Ngo et al, 2014). The prolonged direct physical contact between the growing tip and the filiform apparatus likely allows a direct exchange of signals which could result in modification of the filiform apparatus cell wall structure. CrRLK1L receptors present an extracellular malectin-like domain (Boisson-Dernier et al, 2011), a tandem organisation of two malectin domains with structural similarity to the di-glucose binding malectin protein (Schallus et al, 2008). The malectin di-glucose binding residues are not conserved in the malectin-like domains of ANX1/2 according to structural data (Du et al, 2018; Moussu et al, 2018). However, direct interactions of FER, ANX1/2 and BUPS1/2 malectin-like domains with the pectin building block polygalacturonic acid have been recently reported (Feng et al, 2018; Lin et al, 2018). An extracellular domain anchored to cell wall components and a cytoplasmic kinase domain capable of inducing downstream signalling make FER and the other CrRLK1L proteins a putative link between cell wall status and cellular responses (Verger & Hamant, 2018). Involvement of FER in root mechanosensing provides additional support for this hypothesis (Shih et al, 2014). Therefore, FER and the related receptors HERK1 and ANJ may be fulfilling a cell wall integrity surveillance function in the filiform apparatus, triggering cellular responses upon changes in the composition or mechanical forces registered at this specialised cell wall structure. Future research in this field will undoubtedly provide new views on how these RLKs integrate pollen-derived cues to ensure tight control of fertilisation.

Receptor complexes are a common feature in signal transduction in multiple cellular processes (Burkart & Stahl, 2017; Couto & Zipfel, 2016; Greeff et al, 2012). Our genetic and biochemical results support possible HERK1-LRE-FER/ANJ-LRE-FER heterocomplexes (Fig. 5 and Fig. 6). LRE and related proteins form complexes with RLKs FER, FLS2 and EFR, making them versatile co-receptors that mediate signal perception in multiple processes (Li et al, 2015; Shen et al, 2017). LRE functions in the maternal control of fertilisation and early seed development (Tsukamoto et al, 2010a; Wang et al, 2017), whereas its homolog LLG1 is restricted to vegetative growth and plant-pathogen interactions (Shen et al, 2017). Uncharacterised LLG2 and LLG3 show pollen-specific expression in microarray data and therefore constitute likely candidates as ANX1/2 and BUPS1/2 receptor complex partners to control pollen tube growth. LRE proteins are thought to stabilise their receptor partners in the plasma membrane and to act as direct co-receptors for the extracellular cues sensed by the RLK (Li et al, 2015). As we found that FER localisation in the filiform apparatus is unaltered in *lre-5* plants, with HERK1/ANJ localisation also not affected, our results do not support the role previously reported for LRE as a chaperone for FER localisation in synergid cells (Li et al, 2015). A strict requirement for LRE as a FER chaperone in the synergid cells has also been challenged by a previous report evidencing that the fertility defect in *lre* female gametophytes could be partially rescued by pollen-expressed LRE (Liu et al, 2016). In the absence of synergid-expressed LRE, the authors speculate that sufficient FER is still localised to the filiform apparatus to interact with LRE on the pollen tube plasma membrane, demonstrating a more minor role for LRE intracellular activity in the synergid cells to correctly localise FER (Liu et al, 2016). We hypothesise that LRE could act as co-receptor for FER and HERK1 or ANJ at the filiform apparatus, forming tripartite HERK1-LRE-FER or ANJ-LRE-FER complexes that sense pollen-derived ligands such as RALF peptides or cell wall components. Structural studies of RLK-LRE complexes will shed light on LRE protein functions in membrane heterocomplexes.

Our results indicate that HERK1, ANJ and LRE are not required to generate the ROS-enriched environment in the micropyle of mature ovules under our experimental conditions, while FER is involved in this process (Fig. 4; (Duan et al, 2014)). The role of FER in ROS production has also been characterised in root hairs, where FER activates NADPH oxidase activity via ROPGEF and RAC/ROP GTPase signalling, ensuring root hair growth stability (Duan et al, 2010). Micropylar ROS accumulation prior to pollen tube arrival depends on NADPH oxidase activity and FER, suggesting a similar pathway to root hairs may take place in synergid cells (Duan et al, 2014). This evidence places FER upstream of ROS production, whereas FER, HERK1/ANJ and LRE would function upstream of pollen tube burst. One possible explanation is that FER is a dual regulator in synergid cells, promoting ROS production and regulating pollen tube reception, while HERK1/ANJ and LRE functions are restricted to the latter under our environmental conditions. Kinase-inactive mutants of FER rescue the pollen tube overgrowth defect in *fer* mutants, but cannot restore the sensitivity to exogenous RALF1 in root elongation (Haruta et al, 2018). These recent findings support multiple signal transduction mechanisms for FER in a context-dependent manner (Haruta et al, 2018). It would thus be informative to test whether the kinase-inactive version of FER can restore the ovular ROS production defect in *fer* mutants. The use of genetic ROS reporters expressed in synergid cells and pollen tubes in live imaging experiments would allow us to observe specific changes in ROS production at the different stages of pollen tube perception in ovules, as performed with Ca^2+^ sensors (Denninger et al, 2014; Hamamura et al, 2014; Ngo et al, 2014). ROS production and Ca^2+^ pump activation in plant cells have been linked during plant-pathogen interactions and are thought to take place during gametophyte communication (Bleckmann et al, 2014; Ma & Berkowitz, 2007). Thus, given the dynamic changes in Ca^2+^ during the different stages of pollen tube reception in synergids and pollen, it is likely that ROS production variations also take place in parallel. Studying ROS production profiles during pollen perception in the *fer-4*, *herk1 anj* and *lre-5* backgrounds would provide the resolution required to link these receptors to dynamic ROS regulation during pollen reception. Induction of specific Ca^2+^ signatures in the synergids upon pollen tube arrival is dependent on FER, LRE and NTA (Ngo et al, 2014). Given that NTA relocalisation after pollen reception depends on functional HERK1/ANJ and NTA is involved in modulating Ca^2+^ signatures in the synergids, it is possible that HERK1 and ANJ might also be required for Ca^2+^ signalling during pollen perception.

Downstream signalling after pollen tube reception in the synergid cells likely involves interactions of HERK1, ANJ and FER with cytoplasmic components through their kinase domain. Our results indicate that the kinase activity of HERK1/ANJ is not required for controlling pollen tube rupture (Fig. S4B), as has been reported for FER (Kessler et al, 2015). The *fer-1* pollen tube overgrowth defect could also be rescued with a chimeric protein comprising the FER extracellular domain and the HERK1 kinase domain (Kessler et al, 2015). This implies that the FER and HERK1/ANJ kinase domains are likely redundant in controlling pollen tube burst and may transduce the signal in a similar manner. Testing whether FER-dependent induction of ROS production in the micropyle is also independent of its kinase activity and whether the HERK1/ANJ kinase domains can also substitute for the FER kinase domain in this process would provide insight into how this signalling network is organised.

This study provides evidence for the involvement of multiple CrRLK1L detectors of pollen tube arrival at the female gametophyte, implicating HERK1 and ANJ as co-receptors of FER. The action of multiple CrRLK1L proteins at the filiform apparatus highlights the key relevance of the CrRLK1Ls in controlling reproduction in flowering plants.

## Methods

### Experimental Model and Subject Details

#### Plant material

*Arabidopsis thaliana* T-DNA insertion lines *herk1* (At3g46290; N657488; *herk1-1;* (Guo et al, 2009a)), and *anj* (At5g59700; N654842; *anj-1*) were obtained from the Nottingham Arabidopsis Stock Centre (NASC; (Alonso et al, 2003; Kleinboelting et al, 2012)). T-DNA lines *fer-4* (At3g51550; N69044; (Duan et al, 2014; Haruta et al, 2014)) and *lre-5* (At4g26466; N66102; (Tsukamoto et al, 2010a)) were kindly provided by Prof. Alice Cheung (University of Massachusetts) and Dr. Ravi Palanivelu (University of Arizona), respectively. Accession Col-0 was used as a wild-type control in all experiments. T-DNA lines were confirmed as homozygous for the insertion by genotyping PCRs. The *anj* mutant line was characterised as a knockout of gene expression in this study by RT-PCR.

#### Growth conditions

Seeds were stratified at 4°C for three days. Seeds were sown directly on soil and kept at high humidity for four days until seedlings emerged. The soil mix comprised a 4:1 (v:v) mixture of Levington M3 compost:sand. Plants were grown in walk-in Conviron growth chambers with 22°C continuous temperature, 16 hours per day of ~120 µmols^−1^m^−2^ light and 60% humidity. For selection of transformants, seeds were surface sterilised with chlorine gas, sown onto half-strength Murashige and Skoog medium (MS; (Murashige & Skoog, 1962)), 0.8% (w/v) agar, pH 5.7 (adjusted with KOH), supplemented with the appropriate antibiotic (25 µg/mL of hygromycin B or 50 µg/mL of kanamycin). Seeds on plates were stratified for three days at 4°C and then transferred to a growth chamber (Snijders Scientific) at 22°C, 16 hours per day of ~90 µmols^−1^m^−2^ of light. Basta selection was carried out directly on soil soaked in a 1:1000 dilution of Whippet (150 g/L glufosinate ammonium; AgChem Access Ltd).

### Method Details

#### Phenotyping

To quantify seed production, fully expanded green siliques were placed on double-sided sticky tape, valves were dissected along the replum with No. 5 forceps, exposing the developing seeds. Dissected siliques were kept in a high humidity chamber until photographed to avoid desiccation.

Carpels from self-pollinated or hand-pollinated flowers at stage 16 were selected for aniline blue staining of pollen tubes. Carpels were fixed at least overnight in a 3:1 solution of ethanol:acetic acid, then softened overnight in 8M NaOH, washed four times in water and incubated for three hours in aniline blue staining solution (0.1% (w/v) aniline blue (Fisons Scientific) in 0.1M K_2_PO_4_-KOH buffer, pH 11). Stained carpels were mounted in 50% glycerol, gently squashed onto the microscope slide and then visualised with epifluorescence or confocal microscopy. Aniline blue fluorescence was visualised on a Leica DM6 or Olympus BX51 epifluorescence microscope using a 400 nm LED light source and a filter set with 340-380 nm excitation, emission filter of 425 nm (long pass) and 400 nm dichroic mirror. Confocal images were acquired using a 403.5 nm laser line, 30.7 µm pinhole size and filter set with 405 nm dichroic mirror and 525/50 nm emission filter cube.

Quick callose staining was carried out by incubating freshly dissected tissue samples in a 1000x dilution of SR2200 (Renaissance Chemicals Ltd) in half-strength MS, 5% (w/v) sucrose, pH 5.7. Samples were mounted in the staining solution directly and visualised under an epifluorescence microscope with the same settings as used for aniline blue staining. Callose-enriched structures like pollen tubes and the filiform apparatus of ovules display a strong fluorescence within 10 minutes of incubation. Only structures directly exposed to the SR2200 solution are stained.

To observe the development of the female gametophyte, we used the confocal laser scanning microscopy method as described by Christensen (Christensen et al, 1997). Ovules were dissected from unpollinated carpels, fixed for 2 hours in a 4% (v/v) solution of glutaraldehyde, 12.5mM sodium cacodylate buffer pH 6.9, dehydrated in an ethanol series (20%-100%, 20% intervals, 30 minutes each) and cleared in a benzyl benzoate:benzyl alcohol 2:1 mixture for 2 hours prior to visualisation. Samples were mounted in immersion oil, coverslips sealed with clear nail varnish and visualised with an inverted Nikon A1 confocal microscope. Fluorescence was visualised with 35.8 µm pinhole size, 642.4 nm laser line and filter set of 640 nm dichroic mirror and 595/50 nm emission filter cube. Multiple z-planes were taken and analysed with ImageJ.

Analyses of expression patterns of *HERK1* and *ANJ* used *promoter::reporter* constructs. *promoter::GUS* reporters were analysed by testing β-glucuronidase activity in Col-0 plants from the T1 and T2 generations. Samples were fixed in ice-cold 90% acetone for 20 minutes, then washed for 30 minutes in 50mM NaPO_4_ buffer pH 7.2. Samples were transferred to X-Gluc staining solution (2mM X-Gluc (Melford Laboratories Ltd), 50mM NaPO_4_ buffer pH 7.2, 2mM potassium ferrocyanide, 2mM potassium ferricyanide and 0.2% (v/v) Triton-X), vacuum-infiltrated for 30 minutes and incubated at 37°C for several hours or overnight. Samples were cleared in 75% ethanol and visualised under a light microscope or stereomicroscope. For the *promoter::H2B-TdTomato* reporters, unpollinated ovules were dissected from the carpels and mounted in half-strength MS, 5% (w/v) sucrose, pH 5.7. RFP signal was detected on a Leica DM6 epifluorescence microscope using a 535 nm LED light source and a filter set with 545/25 nm excitation filter, 605/70 nm emission filter and a 565 nm dichroic mirror. DIC images were taken in parallel.

H_2_DCF-DA staining of ROS in ovules was carried out as per (Duan et al, 2014). Ovules from unpollinated carpels were dissected and incubated in staining solution (25µM H_2_DCF-DA (Thermo Scientific), 50mM KCl, 10mM MES buffer pH 6.15) for 15 minutes. Samples were subsequently washed three times in H_2_DCF-DA-free buffer for 5 minutes, mounted on slides and immediately visualised by epifluorescence microscopy. H_2_DCF-DA fluorescence was visualised using a 470 nm LED light source and a filter set with 470/40 nm excitation filter, 460/50 nm emission filter and 495 nm dichroic mirror.

All steps were performed at room temperature unless otherwise specified. Ovules were dissected by placing carpels on double-sided sticky tape, separating the ovary walls from the replum with a 0.3 mm gauge needle, and by splitting the two halves of the ovary along the septum with No. 5 forceps. GFP was visualised by epifluorescence microscopy with the same settings used to visualise H_2_DCF-DA fluorescence. TdTomato was visualised as described above.

#### Cloning and transformation of Arabidopsis

To study the cellular localisation and to complement the pollen overgrowth defect we generated the constructs *pANJ::ANJ-GFP*, *pHERK1::HERK1*, *pFER::FER-GFP, pANJ::ANJ-KD-GFP*, and *pHERK1::HERK1-KD*. Genomic regions of interest (spanning 2 kb upstream of the start codon ATG and the full coding sequence excluding stop codon) were amplified by PCR with Phusion DNA polymerase (NEB). *Promoter::CDS* amplicons were cloned via KpnI/BamHI restriction sites into a pGreen-IIS backbone (Basta resistance; from Detlef Weigel’s group, Max Planck Institute for Developmental Biology; (Mathieu et al, 2007)), with or without an in-frame C-terminal GFP coding sequence. Kinase-dead versions of *HERK1* and *ANJ* were generated by targeted mutagenesis of the activation loop residues D606N/K608R of ANJ and D609N/K611R of HERK1 using *pANJ::ANJ-GFP* and *pHERK1::HERK1* constructs as template (Ho et al, 1989). To generate the GUS and H2B-TdTomato reporter constructs, *pHERK1* and *pANJ* (from 2 kb upstream of the ATG start codon) were cloned with a pENTR-dTOPO system (Thermo Scientific) and then transferred to the GUS expression cassette in the pGWB433 destination vector or pAH/GW:H2B-TdTomato via LR recombination [LR clonase II; Thermo Scientific; (Nakagawa et al, 2007)]. ASE *Agrobacterium tumefaciens* strain was used with pGreen vectors; GV3101pMP90 strain was used otherwise. Arabidopsis stable transformants were generated through the floral dip method.

To test interaction *in vivo* in co-immunoprecipitation assays, we generated *pFER::ANJ-GFP* via three-way ligation cloning of KpnI-*pFER*-NotI and NotI-*ANJ*-BamHI fragments into a pGreen-IIS backbone (Basta resistance; from Detlef Weigel’s group, Max Planck Institute for Developmental Biology; (Mathieu et al, 2007)). To test direct interaction between HERK1exJM, ANJexJM and LRE in yeast, we cloned the extracellular juxtamembrane sequence corresponding to the 81 amino acids N-terminal of the predicted transmembrane domain of HERK1 and ANJ, as well as the sequence corresponding to the amino acids 23-138 of LRE [as per (Li et al, 2015)]. Interaction between HERK1, ANJ and FER was also assayed by Y2H and the extracellular domains excluding the signal peptide (HERK1-ECD, amino acids 24-405; ANJ-ECD, amino acids 25-405; FER-ECD, amino acids 28-446) as well as the cytosolic kinase domains (HERK1-KIN, amino acids 429-830; ANJ-KIN, amino acids 429-830; FER-KIN, amino acids 470-895). Amplicons of exJM and KIN domains were cloned into yeast two hybrid vectors pGADT7 and pGBKT7 via SmaI restriction digests, in frame with the activation or DNA binding domains (AD or BD, respectively). Amplicons of ECD domains were cloned into PCR8 entry vectors and subsequently recombined into pGADT7-GW and pGBKT7-GW via LR recombination. Col-0 genomic DNA was used as the template for all cloning events unless otherwise specified.

To mutate *FER* in the Col-0, *herk1 anj* and *herk1 anj lre* genotypes, CRISPR-Cas9 with two guide RNAs was used to generate large deletions. The guide RNAs were designed with https://crispr.dbcls.jp to target two regions of the *FER* gene 1.7 to 2.2 kb apart and were cloned into pBEE401E. T1 transformants were selected with BASTA and based on a *fer-4*-like phenotype. Seed set was assessed in the T2 generation and the lines genotyped at *FER* to verify either a large deletion in the gene or no amplification due to loss of the primer binding sites. Primers used for cloning are listed in Supplementary Table S2.

#### Genotyping PCRs and RT-PCRs

Genomic DNA was extracted from leaves of 2-week old seedlings by grinding fresh tissue in DNA extraction buffer (200mM Tris-HCl pH 7.5, 250mM NaCl, 25mM EDTA and 0.5% SDS), precipitating DNA with isopropanol, washing pellets with 75% EtOH and resuspending DNA in water. Genotyping PCRs were performed with Taq polymerase and 35 cycles with 60°C annealing temperature and one minute extension time. RNA was extracted with the E.Z.N.A. plant RNA extraction kit (Omega Bio-Tek) from 100 mg of floral tissue from multiple plants per line. RNA concentrations were normalised, an aliquot was DNaseI-treated and subsequently transcribed into first strand cDNA with the RevertAid cDNA synthesis kit (Thermo Scientific) using random hexamers. RT-PCR of *ANJ* and the control gene *FER* were performed with the conditions used in genotyping PCRs with 45 seconds of extension time. Primers for genotyping and RT-PCR are listed in the Supplementary Table S2.

#### Yeast two-hybrid assays

Direct interaction assays in yeast were carried out following the Clontech small-scale LiAc yeast transformation procedure. Yeast strain Y187 was transformed with pGADT7 constructs and yeast strain Y2HGold with pGBKT7 constructs (including empty vectors as controls). Yeast diploids cells carrying both plasmids were obtained by mating and interaction tests were surveyed on selective media lacking leucine, tryptophan and histidine.

#### Co-immunoprecipitation and western blots

For assays using transient expression, leaves of 4.5-week-old *N. benthamiana* were infiltrated with *A. tumefaciens* strain GV3101 carrying constructs indicated in figure captions. In all cases, leaves were co-infiltrated with *A. tumefaciens* carrying a P19 silencing suppressor. Leaves were harvested 2 days post-infiltration and frozen in liquid nitrogen before extraction in buffer (20 mM MES pH 6.3, 100 mM NaCl, 10% glycerol, 2 mM EDTA, 5 mM DTT, supplemented with 1% IGEPAL and protease inhibitors). Immunoprecipitations were performed in the same buffer with 0.5% IGEPAL for 3-4 hours at 4 °C with GFP-trap resin (Chromotek). Beads were washed 3 times with the same buffer and bound proteins were eluted by addition of SDS loading dye and heating to 90°C for 10 min. Proteins were separated by SDS-PAGE and detected via Western blot following blocking (in TBS-0.1% Tween-20 with 5% non-fat milk powder) with the following antibody dilutions in the same blocking solution: α-GFP-HRP (B-2, sc-9996, Santa Cruz), 1:5000; α-HA-HRP (3F10, Roche), 1:3000.

To test whether HERK1 associates with FER *in planta*, T2 generation *herk1 anj* lines expressing *pFER::HERK1-GFP* were germinated on selection for 5 days. Homozygous *p35S::Lti6b-GFP* (Col-0 background) was used a control membrane-localized GFP-tagged protein (Kadota et al, 2014). 5-day-old seedlings were transferred to liquid MS culture and grown in 6-well plates for an additional 7 days. Seedlings were harvested and ground in liquid nitrogen and total protein was extracted in IP buffer (50 mM Tris-Cl pH 7.5, 150 mM NaCl, 2 mM EDTA, 10% glycerol, supplemented with 5 mM DTT, 0.5 mM PMSF, Sigma protease inhibitor cocktail P9599, and Sigma phosphatase inhibitor cocktails 2 and 3) + 1% IGEPAL. Extracts were clarified by centrifugation at 10,000*g*, filtered through Miracloth (Millipore), and diluted with detergent-free IP buffer to 0.5% IGEPAL (final concentration). IPs were performed with GFP-trap resin (Chromotek) for 4h at 4°C with mixing. Beads were collected by centrifugation at 500*g* and washed three times with IP buffer + 0.5% IGEPAL. Bound proteins were eluted by heating to 80°C in 2x SDS-loading dye. FER was detected using anti-FER (rabbit polyclonal, 1:1000; Xiao, Stegmann *et al*., under revision) and anti-Rabbit IgG (whole molecule)–HRP (Sigma A0545, 1:5000).

#### Microscopy and image building

Epifluorescence images were obtained with Leica DM6 or Olympus BX51 widefield microscopes equipped with HC PL Fluotar objectives or UPlanFl 4x,10x and 20x objectives, respectively. A Nikon A1 inverted confocal laser scanning microscope fitted with Plan Fluor 40x oil and Plan Apo VC 60x oil objectives was used to obtain confocal micrographs. A Leica M165 FC stereomicroscope was used to visualise floral tissues from GUS stained samples. Leica LASX, NIS Elements Viewer and ImageJ software were used to analyse microscopy images. Inkscape was used to build all figures in this article.

### Quantification and Statistical Analysis

Leica LASX software was used to obtain relative fluorescence intensity profiles from synergid cells by defining linear regions of interest across the synergid cytoplasm in a micropylar to chalazal orientation. Synergid cytoplasm area was defined between the filiform apparatus and the synergid-egg cell chalazal limit using the corresponding DIC images.

Statistical significance in seed set averages and relative fluorescence averages (at equivalent distances from the filiform apparatus) were assessed with Student’s *t*-tests. χ-square tests were used to compare distributions obtained in pollen tube overgrowth assays and ROS measurements in ovules, using the distribution obtained in wild-type plants as the expected distribution. In all tests, *p<0.05, **p<0.01, and ***p<0.001. Sample size *n* is indicated in the graphs or in figure legends.

## Supporting information

Combined Supplemental Information

## Acknowledgements

We thank: Andrew Fleming and his group at the University of Sheffield for early feedback and guidance on experiments; Alice Cheung and Qiaohong Duan from the University of Massachusetts for advice on the ROS assays and for sharing *fer-4* seeds with us; Chao Li from East China Normal University for the *p35S::HA-LRE* construct; Ravi Palanivelu from the University of Arizona for *lre-5* seeds; Martin Bayer from the Max Planck Institute for Developmental Biology for the *pLAT52::TdTomato* line; Ueli Grossniklaus from the University of Zurich for the *pFER::HERK1-GFP* and *pLRE::LRE-Citrine* constructs; Sharon Kessler from Purdue University for sharing the *pMYB98::NTA-GFP* construct; Daphne Goring from the University of Toronto for the pBEE401E CRISPR/Cas-9 construct and Melinka Butenko at the University of Oslo for the pAH21\GW vector used to make *promoter::H2B-TdTomato* reporters. S. G-T. was supported by a Department of Animal and Plant Sciences postgraduate teaching fellowship. Research in J.E.G.’s lab is supported by RCUK grant BB/N004167/1. T. A. D. was supported by a long-term post-doctoral fellowship from the European Molecular Biology Organisation (LTF 100-2017). N. B-T. was supported by a MINECO FPI Fellowship (BES-2014-068868) and we acknowledge David Alabadi for his supervision of N. B-T. The Zipfel laboratory was supported by the Gatsby Charitable Foundation and European Research Council (PEPTALK).

## Author contributions

Conceptualization, S.G-T. and L.M.S.; Methodology, S.G-T. and L.M.S.; Investigation, S.G-T., N.B-T., T.A.D., L.M.S. and E.S.W.; Writing – Original Draft, S.G-T. and L.M.S.; Writing – Review & Editing, all authors; Supervision, C.Z., J.E.G and L.M.S.

## Declaration of interests

The authors declare no competing interests.

