## Supplementary material for "CrRLK1L receptor-like kinases HERCULES RECEPTOR KINASE 1 and ANJEA are female determinants of pollen tube reception": Combined Supplemental Information

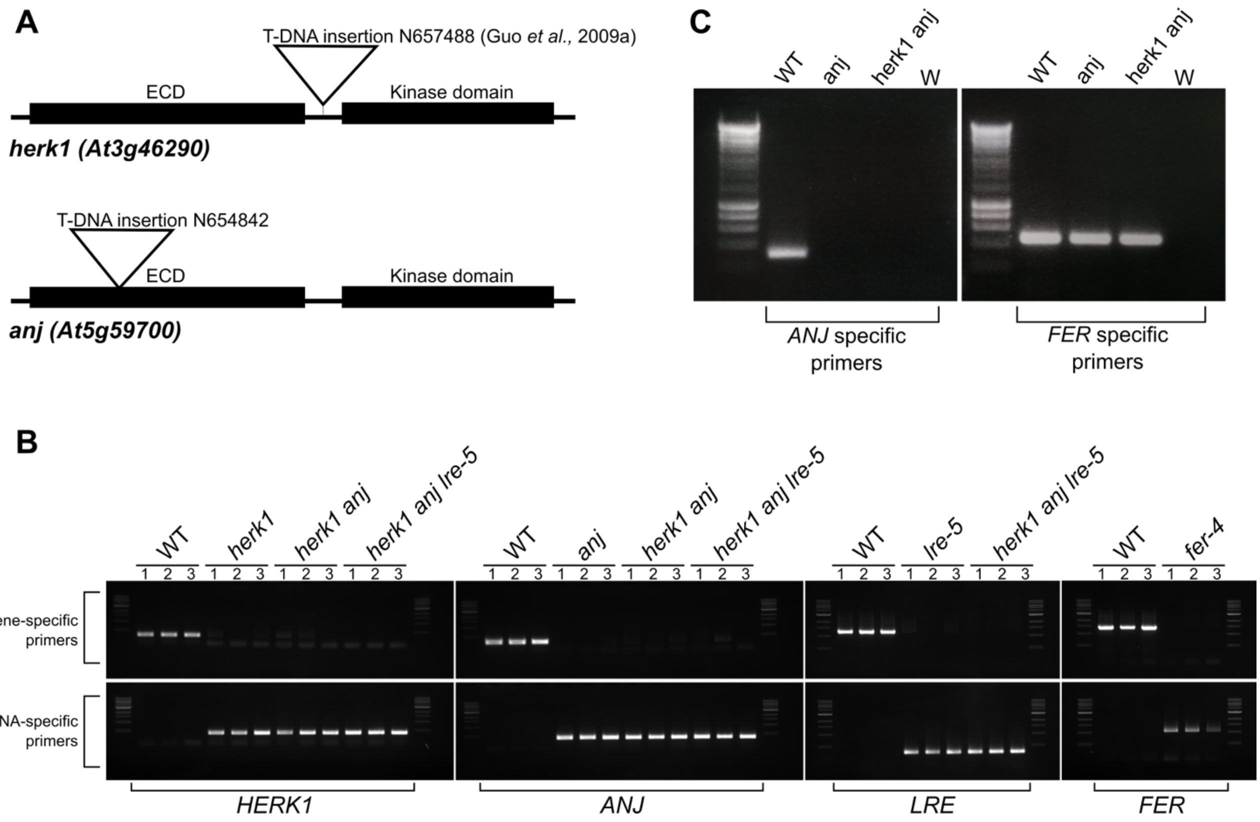


**Figure S1. Confirmation of *ANJEA* gene expression knock out and genotyping of T-DNA lines used in this study.** (A) Diagram of the domain organisation of *HERK1* and *ANJEA* and T-DNA insertion sites in the lines used in this study, *herk1-1* and *anj-1*. (B) Genotyping PCRs to verify homozygosity in the lines used in this study. DNA from three independent seedlings per line was analysed. (C) RT-PCR analysis of *ANJ* gene expression in wild-type, *anj* and *herk1 anj* plants. RNA was extracted from multiple inflorescences from five plants per line. W indicates a water control with no cDNA added to the RT-PCR reaction.

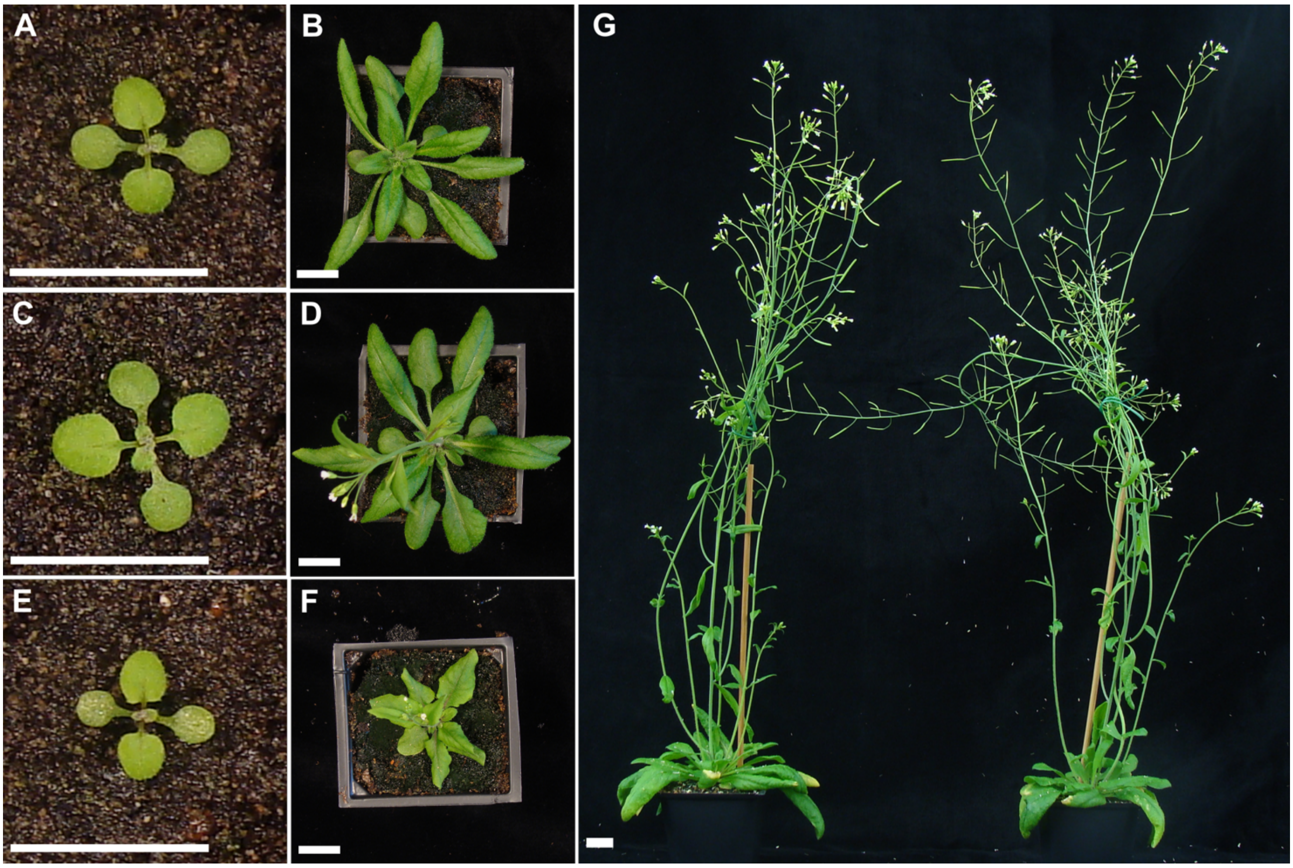


**Figure S2. Growth comparison of WT and *herk1 anj* plants.** (A-B) Representative wild-type plants at 10 and 21 days old. (C-D) Representative *herk1 anj* plants at 10 and 21 days old. (E-F) Representative *fer-4* plants at 10 and 21 days old. (G) Representative wild-type and *herk1 anj* plants (left and right, respectively) at 5 weeks old. Scale bars = 1.5 cm.

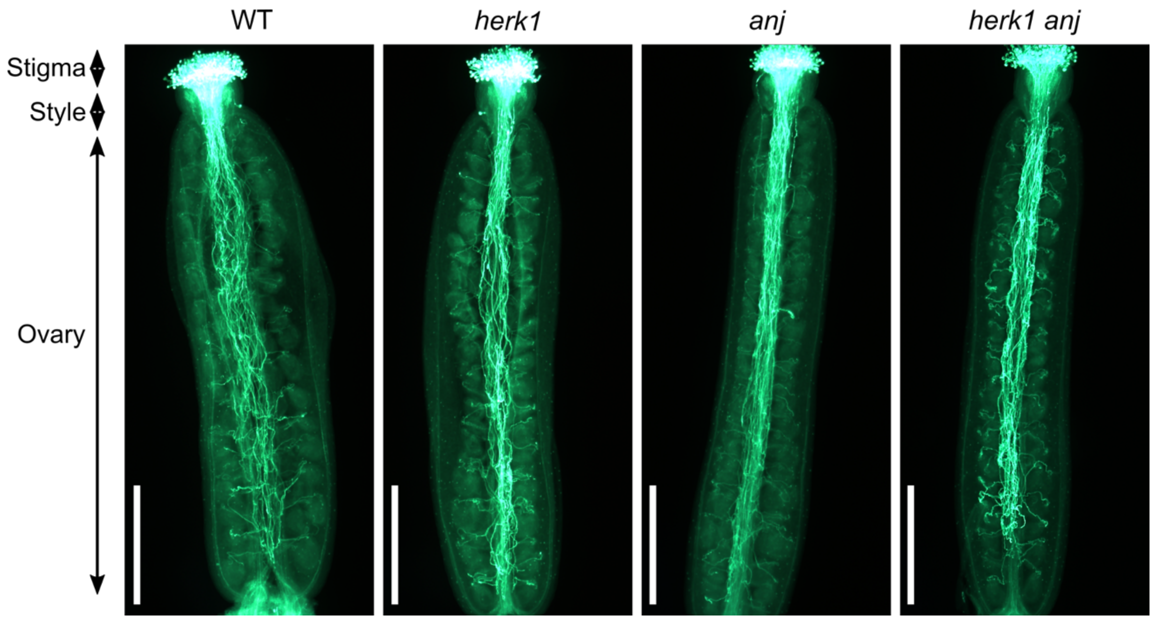


**Figure S3. Pollen tube growth and targeting of ovules is not altered in *herk1 anj* plants.** Aniline blue staining of pollen tubes in self-pollinated stage 16 flowers in wild-type, *herk1*, *anj* and *herk1 anj* plants. Scale bars = 500 μm.

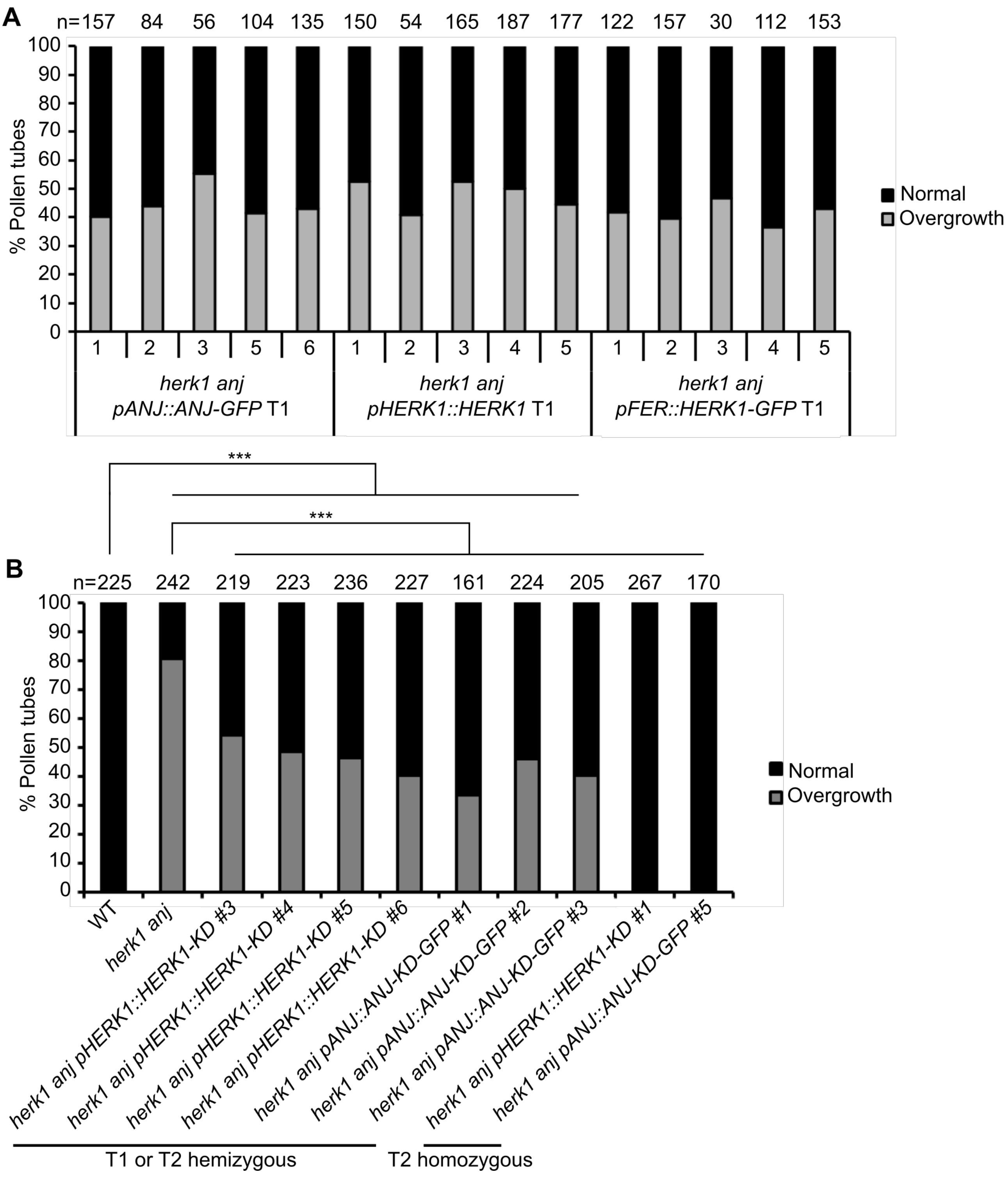


**Figure S4. The *herk1 anj* defect in pollen tube reception can be complemented by expression of *HERK1*, *ANJ*, *HERK1-KD* and *ANJ-KD* constructs.** (A) Percentage of pollen tubes with normal reception at the female gametophyte (black bars) and displaying overgrowth (grey bars) in siliques of five independent T1 *herk1 anj* plants transformed with *pANJ::ANJ-GFP*, *pHERK1::HERK1* and *pHERK1::HERK1-GFP*. Pollen tube reception was scored for ovules in at least three siliques per line. (B) Percentage of pollen tubes with normal reception at the female gametophyte (black bars) and displaying overgrowth (grey bars) in WT, *herk1 anj* plants and at least 4 independent lines of *herk1 anj* transformed with *pHERK1::HERK1-KD* or *pANJ::ANJ-KD-GFP* from generations T1 or T2. Pollen tube reception was scored for ovules in at least three siliques per line. *** $p<0.001$ ($\chi$-square tests).

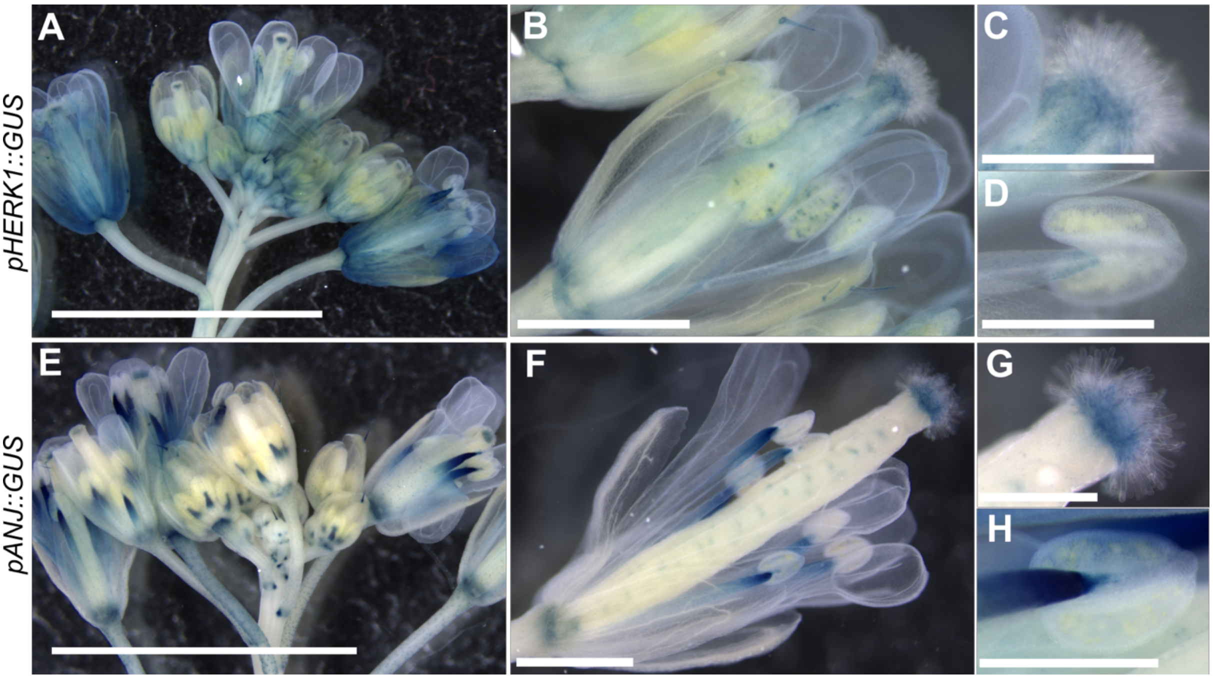


**Figure S5. Expression pattern of *HERK1* and *ANJ* in flowers.** (A-D) Representative image of the expression pattern in inflorescences and flowers of *HERK1* as shown by *pHERK1::GUS*. Details of a mature stigma and anther are shown in (C) and (D), respectively. GUS activity in at least four T1 lines was examined. (E-H) Representative image of the expression pattern in inflorescences and flowers of *ANJ* as shown by *pANJ::GUS*. Details of a mature stigma and anther are shown in (G) and (H), respectively. GUS activity in at least four T1 lines was examined. Scale bars = 5 mm in (A,E) 1 mm in (B,F); 0.5 mm in (C,D,G,H).

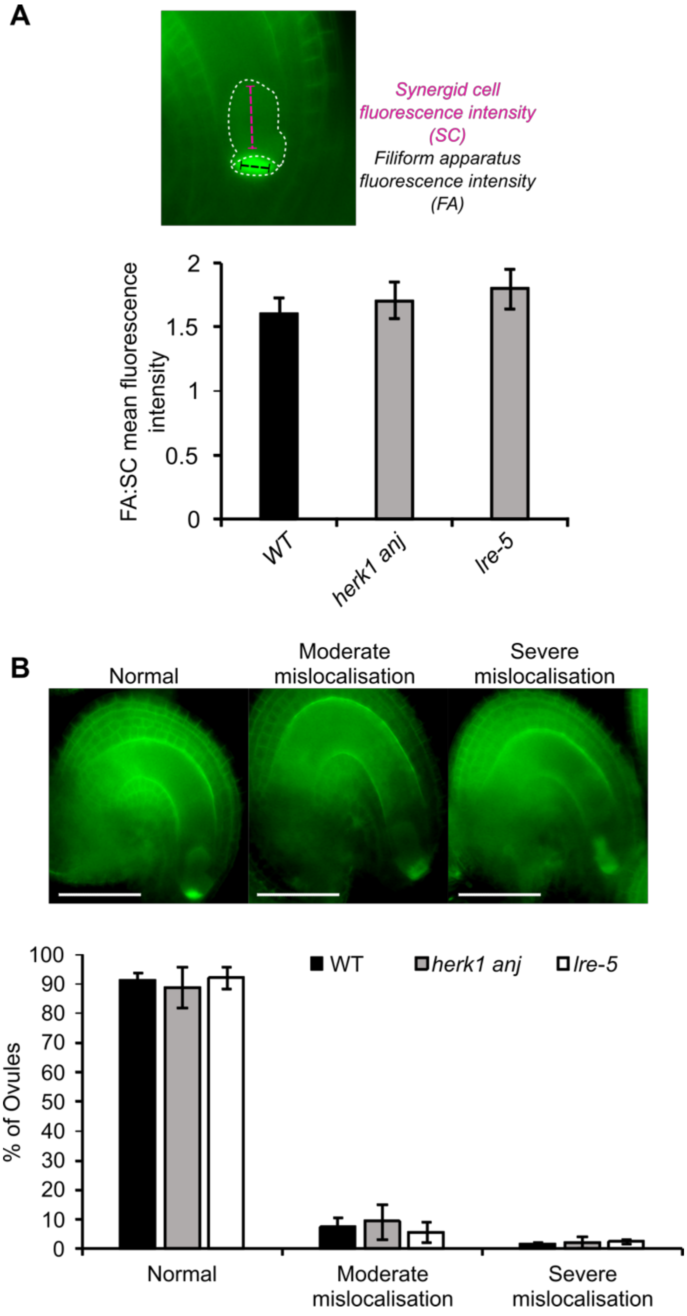


**Figure S6. Quantification of FER-GFP mislocalisation in the synergid cells of *herk1 anj* and *lre-5* ovules.** (A) Ratio between fluorescence intensities at the filiform apparatus (FA) and the synergid cell cytoplasmic region (SC) in mature ovules from wild-type (Col-0), *herk1 anj* and *lre-5* emasculated flowers expressing *pFER::FER-GFP*. Fluorescence profiles for each region of the synergid cells were recorded as exemplified in the upper panel and averaged prior to the ratio calculation (Student's *t* tests, p>0.05). (B) Quantification of moderate and severe mislocalisation defects in the accumulation of FER-GFP at the filiform apparatus in mature ovules from wild-type (Col-0), *herk1 anj* and *lre-5* emasculated flowers expressing *pFER::FER-GFP*. Ovules with clear

FER-GFP expression were assigned to one of the three categories presented in the upper panel, as per (Li et al, 2015). Ovules were obtained from three siliques per plant and three plants per line (total of ovules analysed per line >95). No statistically significant differences were detected in Student's *t* test comparisons with wild-type.

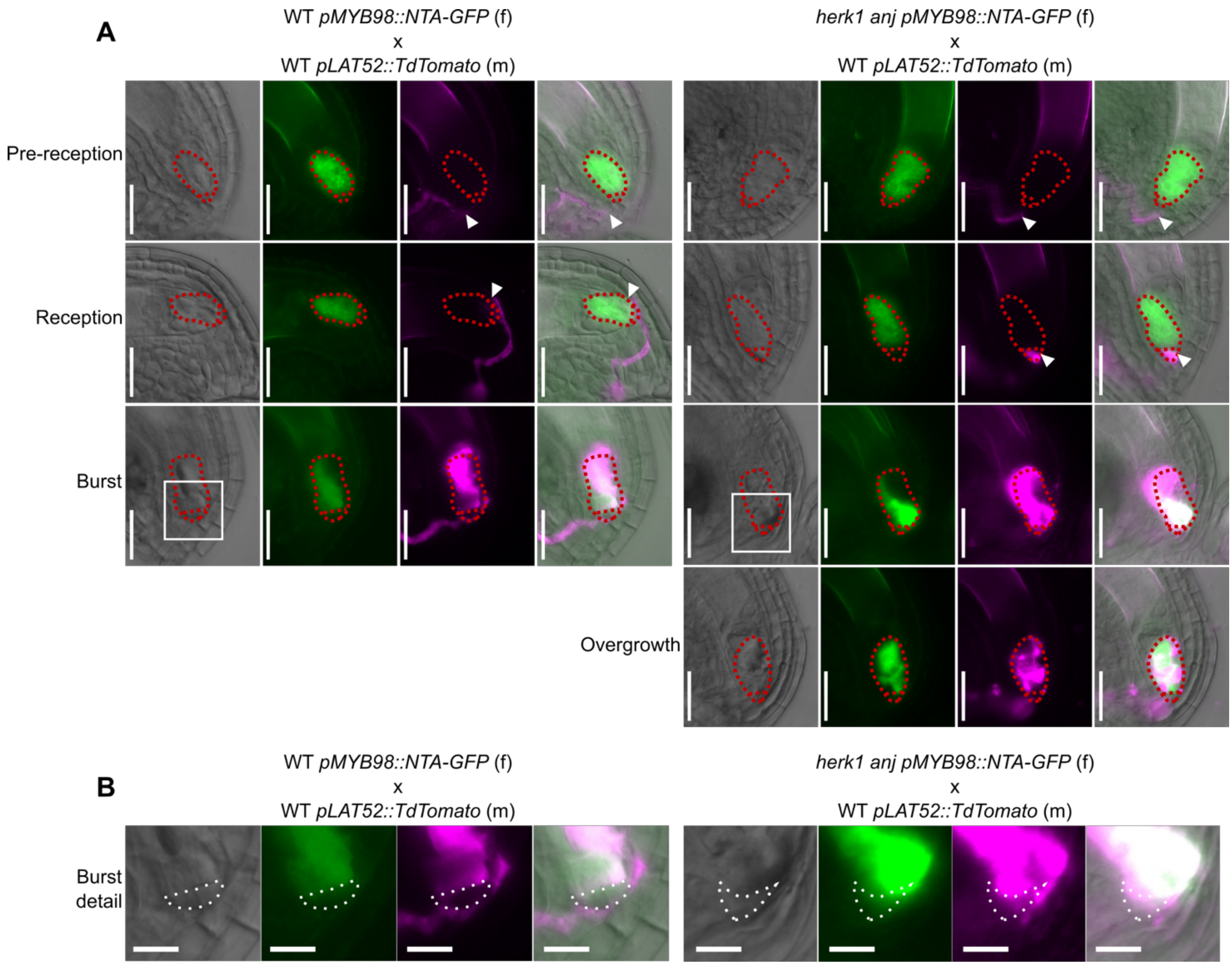


**Figure S7. NTA localisation in the synergid cells of WT and *herk1 anj* at different stages of pollen tube reception.** (A) DIC images are shown in grey. In green is NTA-GFP fluorescence in ovules expressing *pMYB98::NTA-GFP*. In magenta, TdTomato fluorescence from pollen tubes expressing *pLAT52::TdTomato*. On the right are merged DIC and fluorescence images. Red dotted lines delineate the synergid cells. White arrowheads indicate the pollen tube tip. (B) Detailed images of the filiform apparatus corresponding to the areas highlighted with white squares in (A). White dotted lines delineate the filiform apparatus. Scale bars = 25 μm in (A) and 10 μm in (B).

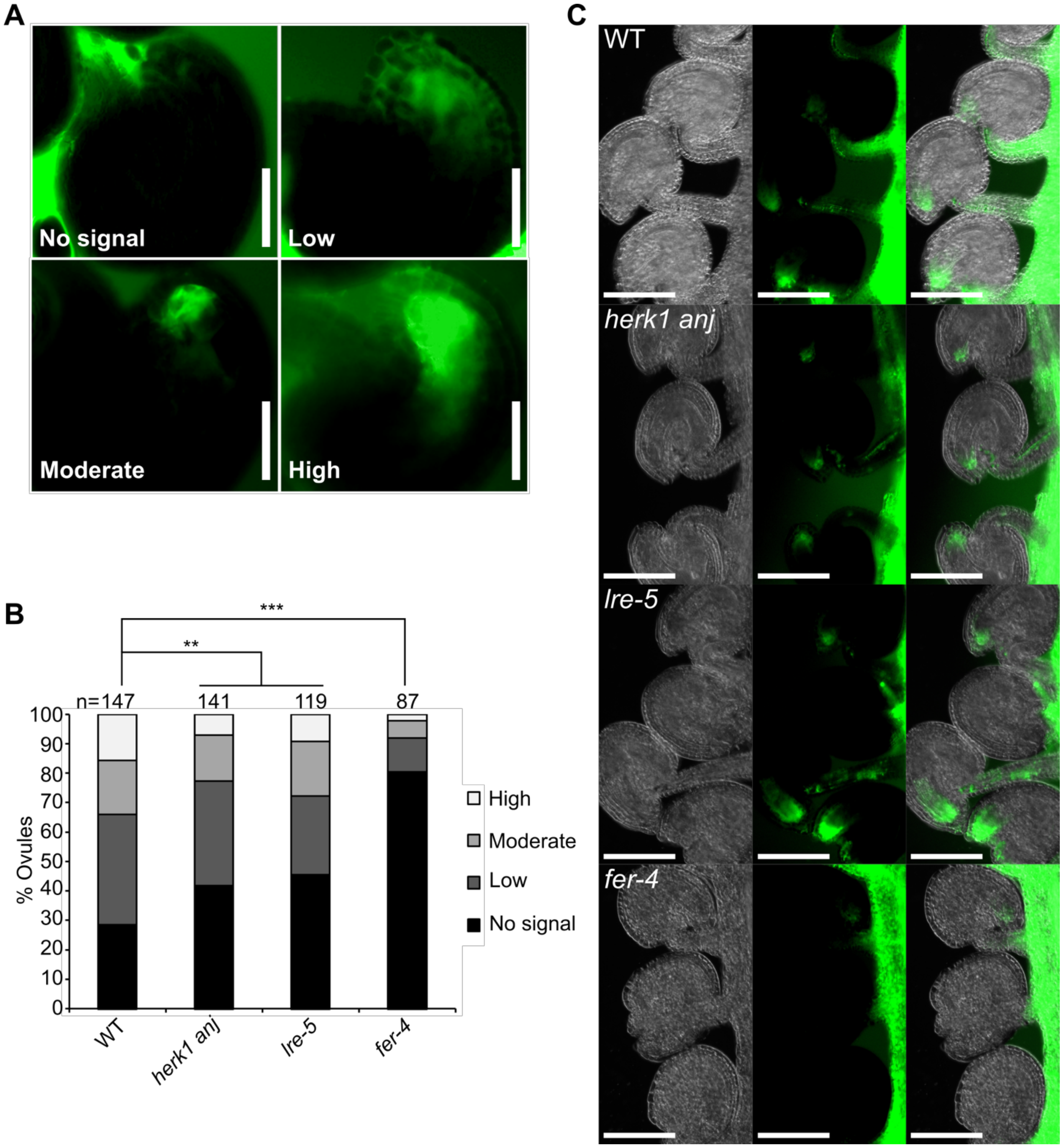


**Figure S8.** **$H_2$DCF-DA measurements of ROS production in *herk1 anj* ovules.** (A) Images of $H_2$CDF-DA fluorescence in representative ovules corresponding to each category used in the ROS assays presented in this study. Scale bars = 25 µm. (B) Quantification of the $H_2$CDF-DA staining of ROS in ovules from wild-type, *herk1 anj*, *lre-5*, and *fer-4* plants at 0 HAE. Categories are listed in the legend. Ovules analysed from six siliques per line. ** $p<0.01$; *** $p<0.001$ ($\chi$-square tests). (C) Representative images of $H_2$CDF-DA staining of ROS in three ovules from wild-type, *herk1 anj*, *lre-5* and *fer-4* plants at 20 hours after emasculation (HAE). Scale bars = 100 µm.

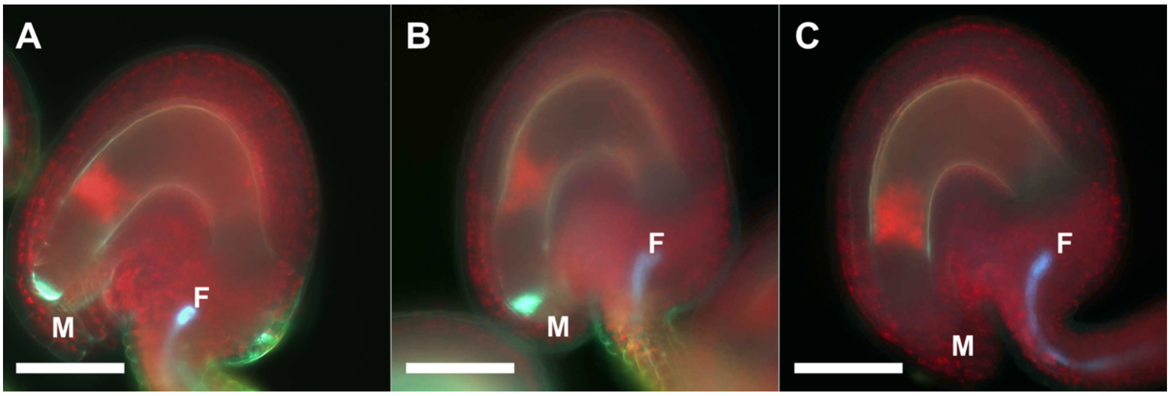


**Figure S9. Callose accumulation at the filiform apparatus in *herk1 anj* mutants.** (A) Representative image of a mature ovule from a wild-type plant. SR2200 white fluorescence at the filiform apparatus indicates accumulation of callose. (B) Representative image of a mature ovule from a *herk1 anj* plant. SR2200 white fluorescence at the filiform apparatus indicates accumulation of callose. (C) Representative image of the background autofluorescence present in mature ovules. Chlorophyll red autofluorescence can be seen in all cell layers in the ovule. Blue autofluorescence from the xylem lignin within the funiculus can also be observed. Scale bars = 25 µm. M, micropyle. F, funiculus.

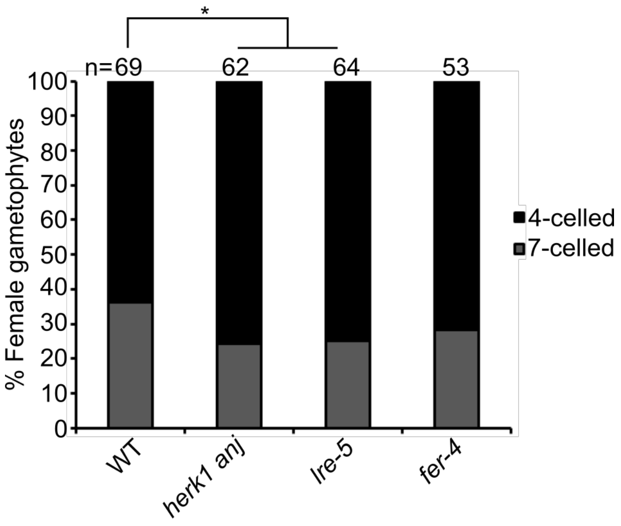


**Figure S10. Female gametophyte development at 20 HAE.** Female gametophyte developmental stage in ovules from stage 14 flowers at 20 hours after emasculation (HAE) in wild-type, *herk1 anj*, *lre-5* and *fer-4* as assessed by confocal microscopy as per (Christensen et al, 1997). Ovules analysed from five siliques per line. * $p<0.05$ ($\chi$-square tests).

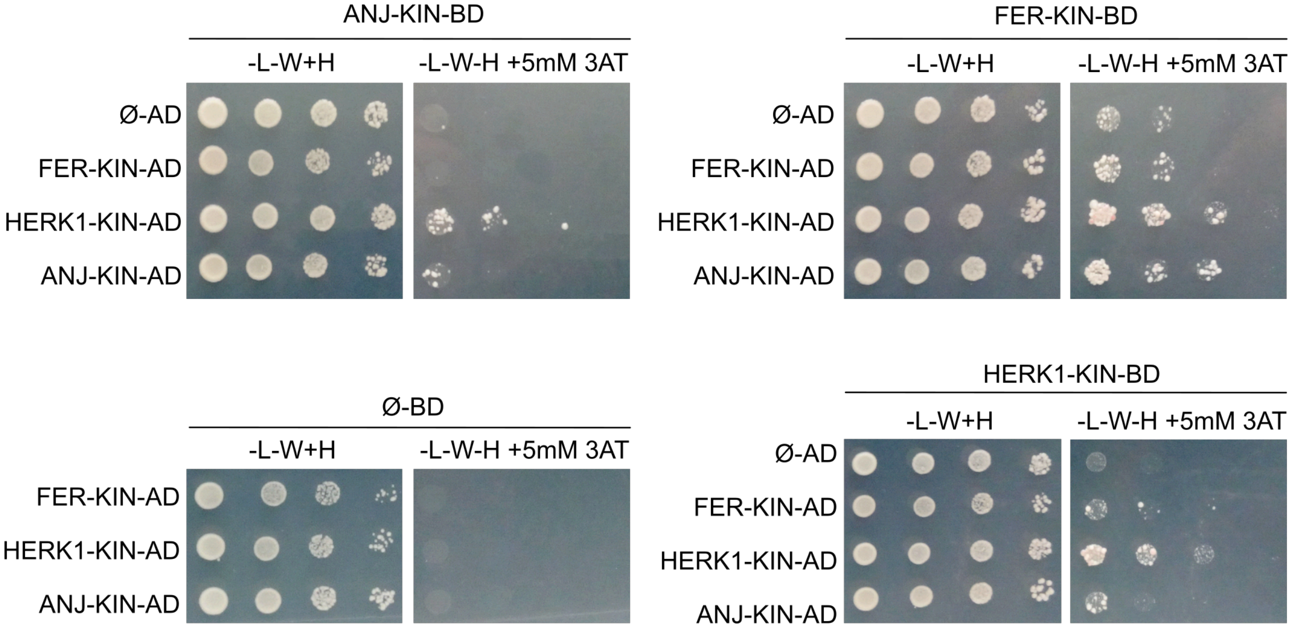


**Figure S11. Yeast two hybrid assays between HERK1, ANJ and FER kinase domains**. Yeast two hybrid assays with the intracellular kinase domains of HERK1, ANJ and FER (HERK1-KIN, ANJ-KIN and FER-KIN, respectively). Ø represents negative controls where no sequence was cloned into the activating domain (AD) or DNA-binding domain (BD) constructs. -L-W-H, growth medium depleted of leucine (-L), tryptophan (-W) and histidine (-H). Plates were supplemented with 5mM 3-Amino-1,2,4-triazole (3 AT) due to yeast growth autoactivation in several of these constructs.

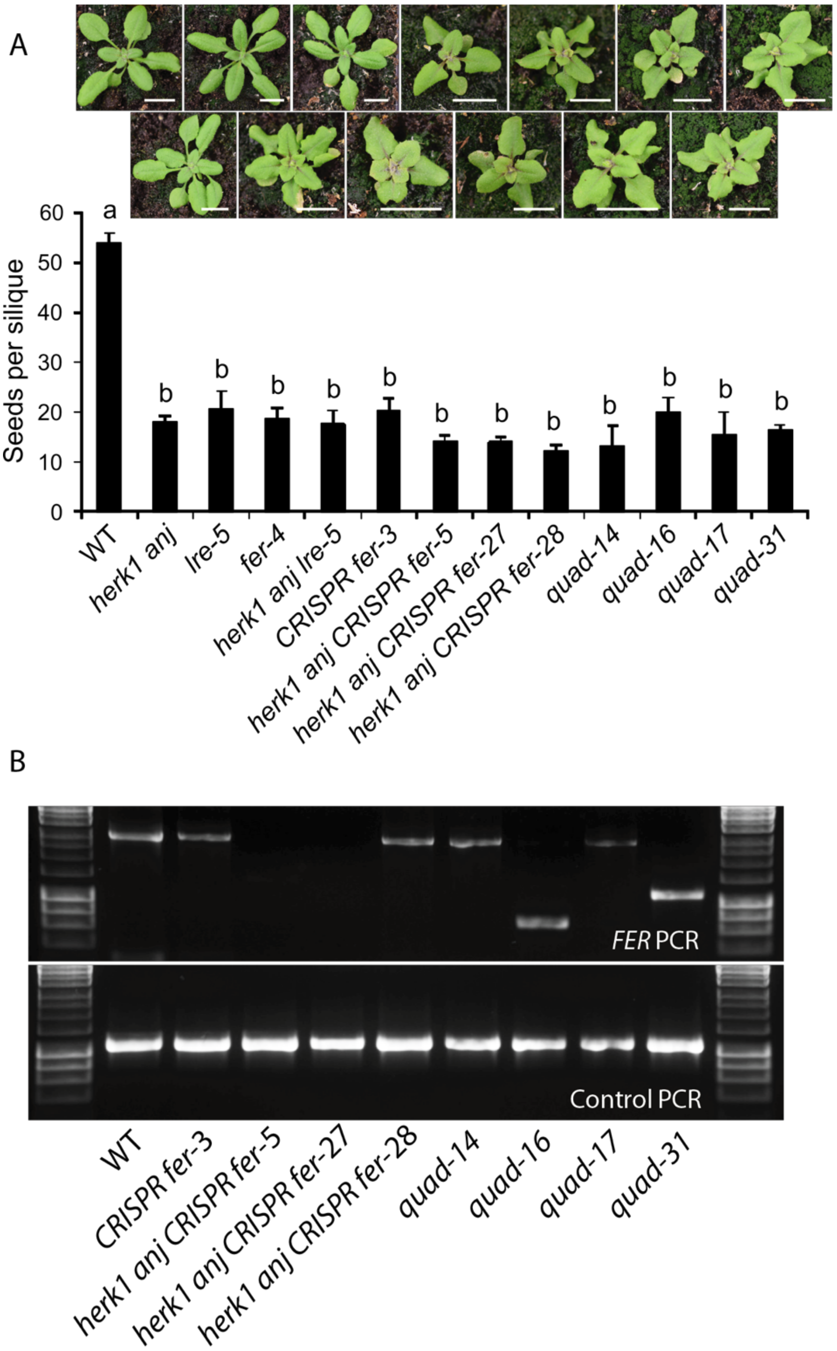


**Figure S12. Quantification of seed set in CRISPR-Cas9 *fer* mutants**. (A) Developing seeds per silique in wild-type, single, double, triple and quadruple mutants as listed. Quad = *herk1 anj lre-5* CRISPR *fer*. Fully expanded siliques were dissected and photographed using an SLR camera. *n* = 15 (three plants per line and five siliques per plant). Data presented are means ± SD. Letters (a, b) mark statistically significant differences between samples in one-way ANOVA analysis followed by Bonferroni's post-hoc comparison of means ($p<0.05$). Pictures above are of plants at 21 days after sowing. Scale bars = 1 cm. (B) PCR amplification of *FER* and control genomic DNA from wild-type and CRISPR-Cas9 *fer* mutants. A lack of amplification from *herk1 anj CRISPR fer* lines 5 and 27 is interpreted as deletion of one or both of the primer binding sites.

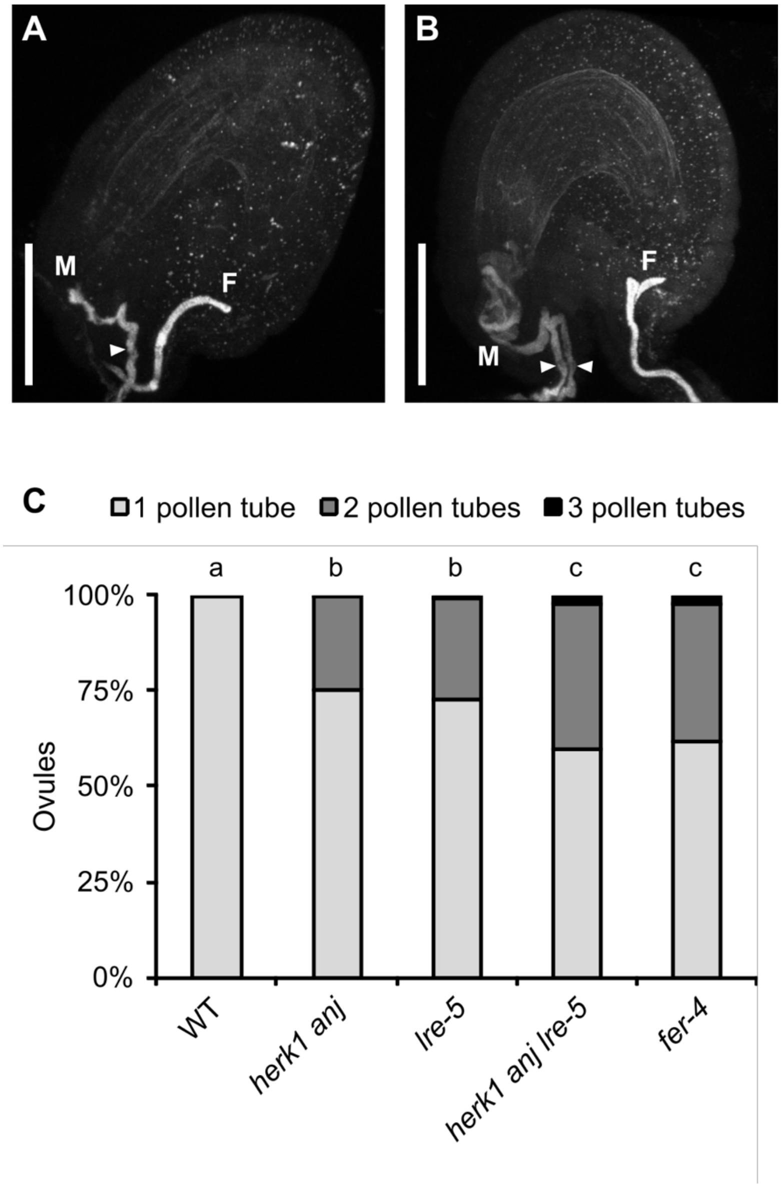


**Figure S13. *herk1 anj* ovules attract multiple pollen tubes.** (A) Representative image of a normal pollen tube reception event in a wild-type ovule by confocal microscopy. (B) Representative image of a *herk1 anj* ovule displaying pollen tube overgrowth and multiple pollen tubes in the micropyle. Images are maximum intensity projections from confocal microscopy images across several z-planes of ovules stained with aniline blue. M, micropyle. F, funiculus. White arrowhead, pollen tube. Scale bars = 50 μm. (C) Polytubey quantification in wild-type (Col-0), *herk1 anj*, *lre-5*, *herk1 anj lre-5* and *fer-4* ovules by epifluorescence microscopy following hand pollination at 24h after emasculation. Ovules from 10 to 13 siliques per line were scored for the number of pollen tubes present at the micropyle if fertilised (total fertilised ovules analysed per line >265). Letters (a, b, c) mark statistically significant differences between samples in multiple Fisher's exact test pairwise comparisons ($p<0.001$).

**Table S1. List of Arabidopsis lines use in this study.** Sources and NASC stock identifiers are listed where relevant.

| **Experimental Models: Organisms/Strains** | | |
|---|---|---|
| *Arabidopsis thaliana:* Col-0 | NASC | N1092 |
| *Arabidopsis thaliana: herk1-1* | NASC | N657488 |
| *Arabidopsis thaliana: anj-1* | NASC | N654842 |
| *Arabidopsis thaliana: fer-4* | Prof. A. Cheung (Duan et al, 2014) | NASC ID: N69044 |
| *Arabidopsis thaliana: lre-5* | Dr. R. Palanivelu (Tsukamoto et al, 2010) | NASC ID: N66102 |
| *Arabidopsis thaliana: herk1-1 anj-1* | This study | N/A |
| *Arabidopsis thaliana: herk1-1 anj-1 lre-5* | This study | N/A |
| *Arabidopsis thaliana:* Col-0 CRISPR *fer* | This study | N/A |
| *Arabidopsis thaliana: herk1-1 anj* CRISPR *fer* | This study | N/A |
| *Arabidopsis thaliana:* Col-0 *herk1 anj lre-5* CRISPR *fer* | This study | N/A |
| *Arabidopsis thaliana:* Col-0 *pHERK1::GUS* | This study | N/A |
| *Arabidopsis thaliana:* Col-0 *pANJ::GUS* | This study | N/A |
| *Arabidopsis thaliana:* Col-0 *pHERK1::H2B-tdTomato* | This study | N/A |
| *Arabidopsis thaliana:* Col-0 *pANJ::H2B-tdTomato* | This study | N/A |
| *Arabidopsis thaliana:* Col-0 *pHERK1::HERK1* | This study | N/A |
| *Arabidopsis thaliana:* Col-0 *pANJ::ANJ-GFP* | This study | N/A |

| | | |
|---|---|---|
| *Arabidopsis thaliana:* Col-0 *pLRE::LRE-Citrine* | This study | N/A |
| *Arabidopsis thaliana:* Col-0 *pMYB98::NTA-GFP* | This study | N/A |
| *Arabidopsis thaliana:* Col-0 *pFER::FER-GFP* | This study | N/A |
| *Arabidopsis thaliana: herk1-1 anj-1 pHERK1::HERK1* | This study | N/A |
| *Arabidopsis thaliana: herk1-1 anj-1 pANJ::ANJ-GFP* | This study | N/A |
| *Arabidopsis thaliana: herk1-1 anj-1 pLRE::LRE-Citrine* | This study | N/A |
| *Arabidopsis thaliana: herk1-1 anj-1 pMYB98::NTA-GFP* | This study | N/A |
| *Arabidopsis thaliana: herk1-1 anj-1 pFER::FER-GFP* | This study | N/A |
| *Arabidopsis thaliana: lre-5 pHERK1::HERK1* | This study | N/A |
| *Arabidopsis thaliana: lre-5 pANJ::ANJ-GFP* | This study | N/A |
| *Arabidopsis thaliana: lre-5 pLRE::LRE-Citrine* | This study | N/A |
| *Arabidopsis thaliana: lre-5 pMYB98::NTA-GFP* | This study | N/A |
| *Arabidopsis thaliana: lre-5 pFER::FER-GFP* | This study | N/A |
| *Arabidopsis thaliana:* Col-0 *pLAT52::TdTomato* | Dr. M. Bayer (unpublished) | N/A |

**Table S2.** **List of primers used for cloning, genotyping PCR or RT-PCR.**

| **Oligonucleotides (5' - 3')** | |
|---|---|
| HERK1 genotyping fw | GTTGCTCGCGGTAGTCTTCT |
| HERK1 genotyping rv | CTGTCCTGAATTCCGCAAGC |
| ANJEA genotyping & RT-PCR fw | CTCCTCTGTAGCAAAACCAGGA |
| ANJEA genotyping & RT-PCR rv | CTCACGTTTACTCCCTCGGG |
| LRE genotyping fw | AAGCCAGTTTTAGAGTACGAAGA |
| LRE genotyping rv | TCAAGTCAACACTAACAAAGCAAAAACAGCGG |
| FER genotyping fw | CGGATCCATGAAGATCACAGAGGGACGATTC |
| FER genotyping rv | CGCAGATCTAGCACCAAACACACAAAACCC |
| FER RT-PCR fw | GAGATGCTCCCTCATTGTACC |
| FER RT-PCR rv | GGCTTACCGCAGACGTAAGC |
| SALK LB genotyping primer | ATTTTGCCGATTTCGGAAC |
| GABI LB genotyping primer | GTGGATTGATGTGATATCTCC |
| pHERK1 fw | TAGGTACCTAGAATGTTTTTCTCAAGTTTTCTTCC |
| HERK1 rv | TAAGGATCCTCTTCCTTCAGATTTCACCAGTTGTG |
| pANJ fw | TTAGGTACCTTGTGGAATCATGAAATCGTAGTGT |
| ANJ rv | TAGGATCCACGTCCCTCAGATTTGATCAGCTGCG |
| pFER fw | TAGGTACCCGAGTTGTAAAAGGCCTGGC |
| FER rv | TAAGGATCCACGTCCCTTTGGATTCATGA |

| | |
|---|---|
| HERK1-KD fw | AGAAACGTGAGATCTGCAAACATATTGCTTGAC GA |
| HERK1-KD rv | AGATCTCACGTTTCTGTGAATGACCGGTTTCGA GT |
| ANJ-KD fw | AGAAACGTCAGATCCGCCAACATATTGCTTGA |
| ANJ-KD rv | GGATCTGACGTTTCTGTGAATCACGGGTTTCG |
| pHERK1 pentrdtopo fw | CACCTAGAATGTTTTTCTCAAGTTTTCTTCC |
| pHERK1 pentrdtopo rv | AACCTGGAAATGGAACAGATC |
| pANJ pentrdtopo fw | CACCTTGTGGAATCATGAAATCGTAGT |
| pANJ pentrdtopo rv | TTCACAAAACCTGGAAATTTTAAATAATT |
| HERK1exJM Y2H (324-406) fw | GGATATTGATCTTAGCACTCTTGTGG |
| HERK1exJM Y2H (324-406) rv | AACCCGAGATTACTCTTACTGCT |
| ANJexJM Y2H (324-406) fw | GCTTGATCTGAGCTCTTATTTATCCA |
| ANJexJM Y2H (324-406) rv | CCACCAACATTCTTCTTAGTGGTTG |
| LRE Y2H (23-138) fw | GATATCGGATGGTGTGTTTGAATCA |
| LRE Y2H (23-138) rv | CCGGCGTTTAGGTTATGTGAATAGAG |
| HERK1 ECD Y2H (24-405) fw | GGATTCACACCTGTGGATAATTAC |
| HERK1 ECD Y2H (24-405) rv | TTACCCGAGATTACTCTTACTGCT |
| ANJ ECD Y2H (25-405) fw | TACGTACCAGTGGATAATTACCTC |
| ANJ ECD (25-405) rv | TTAACCAACATTCTTCTTAGTGGTTG |
| FER ECD Y2H (28-446) fw | GCTGATTACTCTCCAACAGAGA |
| FER ECD Y2H (28-446) rv | TTACGTATTGCTTTTCGATTTCCTAG |
| HERK1 kin Y2H (429-830) fw | GAAGAAGCGGAAACGTGGC |

| | |
|---|---|
| HERK1 kin Y2H (429-830) rv | CCTCTTCCTTCAGATTTCACCAGTTGTG |
| ANJ kin Y2H (429-830) fw | GAAGAAACGAGGACGAGACC |
| ANJ kin Y2H (429-830) rv | CCTCCACGTCCCTCAGATTTGATCAGCTGCG |
| FER kin Y2H (470-895) fw | GGCTTACCGCAGACGTAAGC |
| FER kin Y2H (470-895) rv | CCACGTCCCTTTGGATTCATGA |
| FER CRISPR construct 1 Out fw<br><br>(5' gRNA 1; target FER sequence underlined) | ATATATGGTCTCGATTGTTCTACCCAAACTCGT<br>ACGAGTT |
| FER CRISPR construct 1 In fw<br>(5' gRNA 1) | TGTTCTACCCAAACTCGTACGAGTTTTAGAGCT<br>AGAAATAGC |
| FER CRISPR construct 1 In rv<br>(3' gRNA 1) | AACCGAGTCCGTCACATTCCCTTCAATCTCTTA<br>GTCGACTCTAC |
| FER CRISPR construct 1 Out rv<br>(3' gRNA 1) | ATTATTGGTCTCGAAACCGAGTCCGTCACATTC<br>CCTTCAA |
| FER CRISPR construct 2 Out fw<br><br>(5' gRNA 2) | ATATATGGTCTCGATTGAAAAGGAGTATGCGGT<br>GACAGTT |
| FER CRISPR construct 2 In fw<br>(5' gRNA 2) | TGAAAAGGAGTATGCGGTGACAGTTTTAGAGC<br>TAGAAATAGC |
| FER CRISPR construct 2 In rv<br>(3' gRNA 2) | AACCGGAAGGCGAGATATCATTCCAATCTCTTA<br>GTCGACTCTAC |
| FER CRISPR construct 2 Out rv<br>(3' gRNA 2) | ATTATTGGTCTCGAAACCGGAAGGCGAGATAT<br>CATTCCAA |
| CRISPR-Cas9 *fer* mutant genotyping fw | CATTGACGCGATTCATGTTT |
| CRISPR-Cas9 *fer* mutant genotyping fw | GATGAAGATCACAGAGGGACG |

| | |
|---|---|
| Control gDNA region for genotyping fw | CTGCCTTACGAGCATTGGTT |
| Control gDNA region for genotyping rv | TAACGCTTCCCAAGGTGATT |

**Table S3.** **List of plasmids used in the present study and their corresponding sources.**

| **Recombinant DNA** | **Reference** |
|---|---|
| *pHERK1::HERK1* in pGreen-IIS | This study |
| *pANJ::ANJ-GFP* in pGreen-IIS | This study |
| *pFER::FER-GFP* in pGreen-IIS | This study |
| *pHERK1:HERK1-KD* in pGreen-IIS | This study |
| *pANJ::ANJ-KD-GFP* in pGreen-IIS | This study |
| *pHERK1::GUS* in pGWB433 | This study |
| *pANJ::GUS* in pGWB433 | This study |
| *pHERK1::H2B-tdTomato in pAH21* | This study |
| *pANJ::H2B-tdTomato in pAH21* | This study |
| *pFER::HERK1-GFP* in pMDC111 | Prof. U. Grossniklaus (Kessler et al, 2015) |
| *pMYB98::NTA-GFP* in pMDC83 | Dr. S. Kessler (Davis et al, 2017) |
| *p35S::HA-LRE* in pSK | Dr. C. Li (Li et al, 2015) |
| *p35S::HA-LRE* in pMLBart | This study |
| *pLRE::LRE-Citrine* in pMDC99 | Prof. U. Grossniklaus (Lindner et al, 2015) |
| *pU6-26::FER 5' gRNA 1; pU6-29::FER 3' gRNA 1* | This study |
| *pU6-26::FER 5' gRNA 2; pU6-29::FER 3' gRNA 2* | This study |
| *pGreen-IIS – Cterm GFP* | (Mathieu et al, 2007) |
| *pGWB433* | (Nakagawa et al, 2007) |

| | |
|---|---|
| *pGADT7* | Clontech |
| *pGBKT7* | Clontech |
| *pAH21\GW* | Dr. M. Butenko |
| *pBEE401E* | Prof. D. Goring (Wang et al, 2015) |
| *pCBC-DT1T2* | Prof. D. Goring (Wang et al, 2015) |
